# Evolutionary refinement of mitochondrial and plastid targeting sequences coincides with the late diversification of land plants

**DOI:** 10.1101/2025.03.13.643027

**Authors:** Parth K. Raval, Carolina García García, Maria-Darline Somoano Sanchez, Sven B. Gould

**Affiliations:** Institute for Molecular Evolution, Heinrich-Heine-University Düsseldorf, 40225 Düsseldorf, Germany

**Keywords:** Protein sorting, organelles, terrestrialization, plastid, mitochondria

## Abstract

Plastids and mitochondria are key to plant survival and adaptation. The evolutionary progress of land plants (embryophytes) witnessed gene and genome duplications, and the expansion of organelle-localised proteins. To deal with the increase of nuclear-encoded proteins, targeting to and import by the mitochondrion and plastid is known to have adapted in multiple ways. It included the addition of entire new import channels and lineage-specific import receptors. Through comparative genomics and experimental biology, we uncover further changes of the organelle import machineries. Their evolution likely served to enhance the rate of protein import and improved its physiological regulation, e.g. via interactions between the import channel and respiratory complex. On the cargo side, nuclear-encoded N-terminal targeting sequences of mitochondrial (mNTS) and plastidal (pNTS) proteins diverged in their charge via a preference for phosphorylatable amino acids (adding negative charges after phosphorylation) and an avoidance of positive charges in the pNTS, which is most evident in eudicots. Using *Chlamydomonas* and *Marchantia*, we experimentally underscore that the evolved NTS divergence prevents mis-sorting between mitochondria and plastids. In line with the increase of phosphorylatable amino acids in the pNTS, we pinpoint the embryophytic origin of a membrane-anchored phosphatase, PAP2, that is associated with targeting sequence processing. On the whole, we propose a revised model for plant organelle protein import evolution from algae to angiosperms, which facilitated flourishing of this lineage on the land.

## Introduction

Mitochondria and plastids, two key compartments of photosynthetic eukaryotes, originate from Gram-negative bacteria^1–3^ and part of their transformation from a free-living organism to an organelle includes the transfer of genetic information to the host nucleus^4^. A plethora of core functions they perform^5–9^ hence hinges upon cytosolic protein biosynthesis and the subsequent import of proteins across a pair of membranes^10–14^. Protein import complexes, consisting of dozens of subunits, serve as a necessary gateway to all organelle functions and by that, they affect fitness and macroevolutionary trends^15,16^. Protein translocation across organelle membranes comprises an interplay of targeting sequences, chaperones, transport channels, and also kinases and phosphatases^14,17,18^. Owing to the challenge of simultaneous protein sorting between two organelles of endosymbiotic origin – that do so based on quite similar principles^15,19,20^ – this process is necessarily more involved in algae and plants and likely has been under a special selection pressure since plastid origin.

The emergence of major archaeplastidal lineages such as the Chloroplastida or Embryophyta witnessed the expansion of organelle proteomes, which likely improved the ability to responds to abiotic stresses^15,21–23^. Translocation across the correct organellar membranes is a key for these proteins to be able to manifest their fitness advantage. The main import channel of the outer chloroplast membrane of the Chloroplastida, TOC75, originated through the duplication of the original import channel OEP80 and allowed for a division of labour between the two^15^. Concomitant with the origin of TOC75 in the green lineage, was the loss of an N-terminal, single amino acid-based targeting motif^15,24,25^, which can discriminate between mitochondrial and plastid cargo in rhodophyte and glaucophyte algae ^26–28^. This event likely affected the quality of protein targeting negatively early on and selected for compensatory mechanisms to evolve. The expansion of organelle proteomes in land plants occurred around the same time^23^, maybe as a consequence, but also providing a positive feedback loop for the selection of a more flexible, yet also accurate cargo recognition.

Several strategies ensure correct protein sorting in the land plant models studied, of which N-terminal targeting sequences (NTS) are one. Mitochondrial NTSs (mNTS) commonly contain an alpha-helical motif, whereas plastidal ones (pNTS) encode a beta sheet ^29,30^. Differences in secondary structure, however, are not yet very well defined and other physio-chemical properties are important, too. In *Arabidopsis*, the first 20 amino acids of the mitochondrial targeting sequence are characterized by a higher positive charge, which is typical for nuclear-encoded mitochondrial matrix proteins in general, whereas those of the pNTS are enriched in serine residues ^31–34^. Phosphorylation of pNTS residues increases the import rate into plastids ^35^ and simultaneously supresses import by mitochondria of otherwise plastid-targeted proteins ^19^. In-silico modelling suggests that features such as the overall charge are sufficient to separate an mNTS from a pNTS^19,32^, but for which more experimental support is needed ^31,36,37^.

Current data regarding the interplay of targeting sequences and import components of mitochondria and plastids is largely restricted to the model system *Arabidopsis* and a few other angiosperms. A comprehensive picture of how conserved the interplay is across Archaeplastida or how changes might have influenced major transitions in algae and plant evolution is missing^38^. We do not fully understand how plastid proteins are exactly distinguished from mitochondrial ones, how dual targeted cargo is selected, or what exact regulatory role the evident cytosolic phosphorylation and de-phosphorylation at the outer membranes of the organelles might play^39–43^. Here, combining comparative genomics, phylogenetics and experimental biology in the chlorophyte *Chlamydomonas* and an early diverging land plant, the bryophyte *Marchantia*, we track the origins and evolution of the protein import components across the Archaeplastida lineage, from algae to angiosperms, and characterize changes that improved accuracy, and likely the rate, of organelle protein import.

## Results

### Divergence of mitochondrial and plastid targeting sequences upon terrestrialization

Membrane receptors and channels can recognize a majority of their cargo based on N-terminal targeting sequences (NTS). To investigate changes in NTSs from algae to angiosperms, we first analyzed the NTS of proteins inferred (based in orthology with experimental proteomes) to be plastid, mitochondrial and dual targeted (pNTS, mNTS, dNTS, respectively) from five major chloroplastidal lineages (Table S1B, Fig. S1). In algae, features of the mNTS and pNTS were rather indistinguishable, suggesting only a weak, if any, selection pressure on the parameters analysed (Fig. 1A-B). Land plants differ and showed converging values across species with a clear separation between pNTSs and mNTSs on two counts: the pNTSs are enriched in phosphorylatable amino acids (Ser, Thr) from streptophyte algae to eudicots (Fig. 1A) and the mNTSs are enriched with respect to a positive charge across the sequences analysed (Fig. 1B). Angiosperms, both mono- and eudicots, showed the clearest separation. The values of NTSs of proteins imported by plastids and mitochondria ranges in between those with only a single destination (Fig. 1A-B), underscoring a compromise with respect to dually targeted proteins.

**Figure 1:**
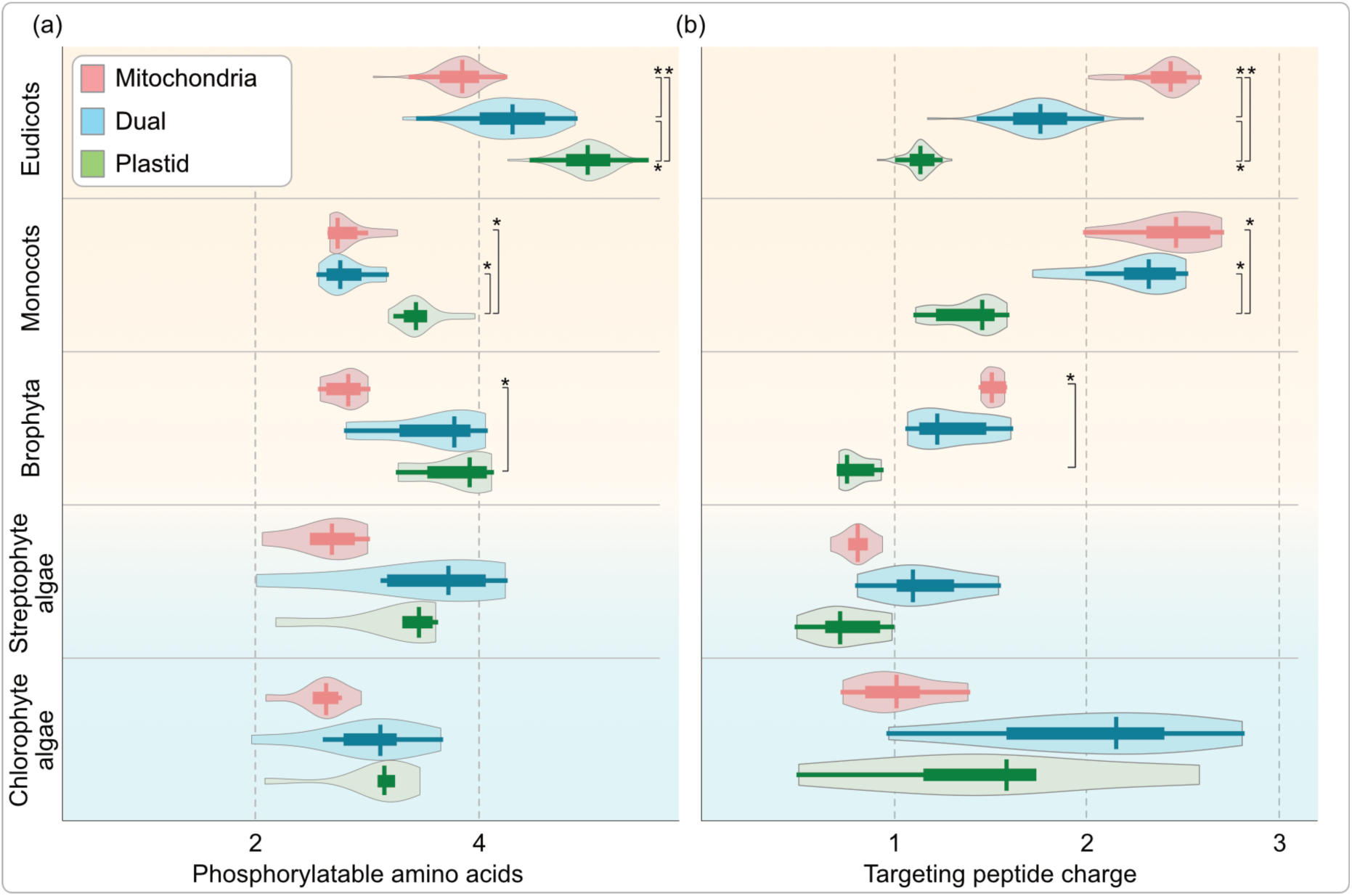
The evolution of amino acid charge across the targeting peptides of photosynthetic eukaryotes. The total number of **(a)** phosphorylatable amino acids (serine+threonine) and **(b)** electric charge for each protein, in a given species, was calculated from the first 20 amino acids of N terminal targeting sequence (NTS) of mitochondrial, plastid and dual targeted proteins. Mean of all proteins of a given organelle in a given species was treated as a representative and plotted for each clade within a violin plot. Each violin therefore represents organelle proteomes of an entire clade, with the top three representing land plants and the bottom two representing algae. Within each violin plot, the vertical line represents the median, the thick horizontal line the interquartile range (from 25th to 75th percentile), and the trailing thin horizontal line representing the full data range. Multiple ANOVA (with Bonferroni correction for multiple comparisons) was performed across and within clades, and organelles. Select statistically significant pairs are shown on the right side (asterisks), full statistics are summarised in the figure source data. N, number of organelle NTSs for a given species from which the mean was calculated to be included in clade-violin plot = 21-2349; n, number of species in a given clade violin = 6-66, see the figure source data for exact numbers.

To investigate where the NTS began to diverge during the evolution of Archaeplastida, we reconstructed the ancestral states of mean charges and phosphorylatable amino acids in NTSs across five plant lineages (Fig. 2, Fig. S2-3). A comparison between eudicots and chlorophyte algae underscores that at the eudicot ancestral node, the pNTS was already enriched in phosphorylatable amino acids and the mNTS accumulated positive charged amino acids, traits much less prominent in the chlorophyte ancestor (Fig. 2A). Within eudicots, the mustard and brassicaceae families showed the highest values and changes in the two traits happened concurrently, i.e. species with a higher positive charge in mNTSs also showed a higher number of phosphorylatable amino acids in the pNTS, and *vice versa*. On the contrary, in chlorophyte algae, changes in these two traits did not occur simultaneously and across species the variation is high (Fig. 2B). It is likely that a selection for the NTSs to diverge increased in embryophytes and in particular in some clades of angiosperms. Generally, phosphorylatable amino acids in pNTSs and positive charges in mNTSs started to increase with the transition from water to land (Fig. 2C-D). Ancestral states of these groups suggest that the divergence of charge predates the diversification within eudicots and angiosperms, and it might have played a role in the evolutionary success of the groups.

**Figure 2:**
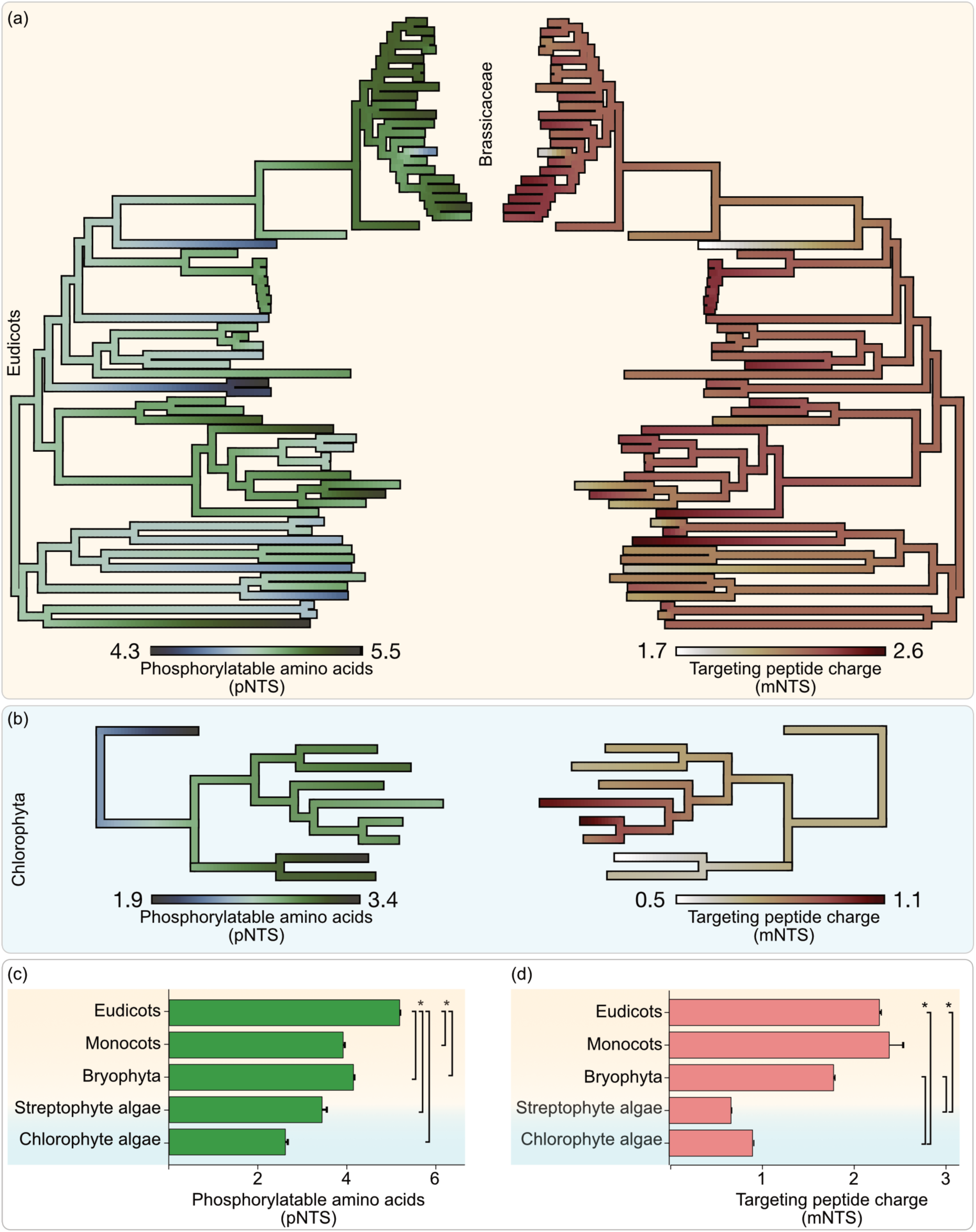
Ancestral state reconstruction of NTS divergence. Ancestral states of phosphorylatable amino acids in pNTSs and charges of mNTSs for five plant lineage was reconstructed using phylogenies for each lineage (Fig. S2) and the phylogenies were color-coded based on these values across ancestors. A complete phylogeny and ancestor state values are shown for **(a)** eudicots and **(b)** chlorophyte algae. Based on a similar reconstruction, inferred means of phosphorylatable amino acid residues across **(c)** all pNTSs and means of positive charges across **(d)** all mNTSs of the last common ancestors of each group is plotted (c-d, *P<0.05, Welch’s t-test). The full ASR of streptophyte algae, bryophytes and monocots, together with the raw data for all five clades, is summarized in Fig. S3 (and its source data).

If NTS divergence is indeed aiding the correct targeting and maintenance of organelle biology, it is expected to be enriched based on protein function and a contribution to defining the organelle’s biology. To test this, we analysed a more reliable localisation and functional annotation dataset of *Arabidopsis thaliana.* We found that plastid proteins involved in photosystem biology and redox reactions are more enriched in phosphorylatable amino acids (Fig. S4) and have a lower positive charge than the rest of plastid proteins and of mNTSs, i.e. the pNTSs of these functional categories are more diverged from the mNTSs than generally observed. A subset of plastid proteins, including the proteins involved in metabolism and translation, have a pNTS charge closer to the mean mNTS charge, i.e. the pNTS of these proteins are less diverged from mNTSs. Some of the key mitochondrial proteins (e.g. TCA cycle proteins) also show a higher mNTS charge than average of all mNTS (Fig. S4), suggesting that these proteins are under a stronger selection pressure to remain specifically mitochondria targeted. Proteins involved in translation in general have a wider distribution of charge and number of phosphorylatable amino acids. It is likely that proteins whose mistargeting is detrimental (e.g. photosystem, redox reactions) are under a stronger selection to diverge from mNTS than the ones whose misstarting is not (e.g. protein synthesis). The latter, over time, provides ground to evolve dual targeting.

In summary, the distribution of NTS features such as charge and phosphorylatable amino acid residues (Fig. 1), their ancestral states (Fig. 2), and the more pronounced feature diversification with respect to the functional category of a protein (Fig. 2, Fig. S4), suggests that these patterns are the result of a strong selection that amplifies targeting sequence divergence, specifically after the origin of land plants. Considering the otherwise general lack of strong differences between the secondary features of pNTS and mNTS, this divergence in the first 20 amino acids emphasizes its important role in the differential recognition of mitochondrial and plastid cargo.

### The divergence of targeting sequences enhances accuracy of protein sorting

To test the hypothesis that the divergence of NTS charge alone has functional consequences for protein targeting, we generated variants of the mitochondrial malic enzyme 1 (MME1) in algal and land plant model systems (Fig. S5). The MME1 homologue of *Chlamydomomas reinhardtii* (*Cr*MME1; Cre06.g268750) has a charge of +5 in the first 20 amino acids and these native 20 amino acids alone are sufficient for correct mitochondrial targeting (Fig. 3). The positive charge is crucial, as constructs in which the overall charge is changed to −5 or 0 fail to be imported by mitochondria. This is in line with an evolutionary early dependence of positive charges for mitochondrial matrix targeting, which manifested itself during eukaryogenesis and in early diverging eukaryotes^44–46^. It was a necessity already during plastid origin and plastid targeting sequences evolved accordingly^47^ (Fig.1-2).

**Figure 3:**
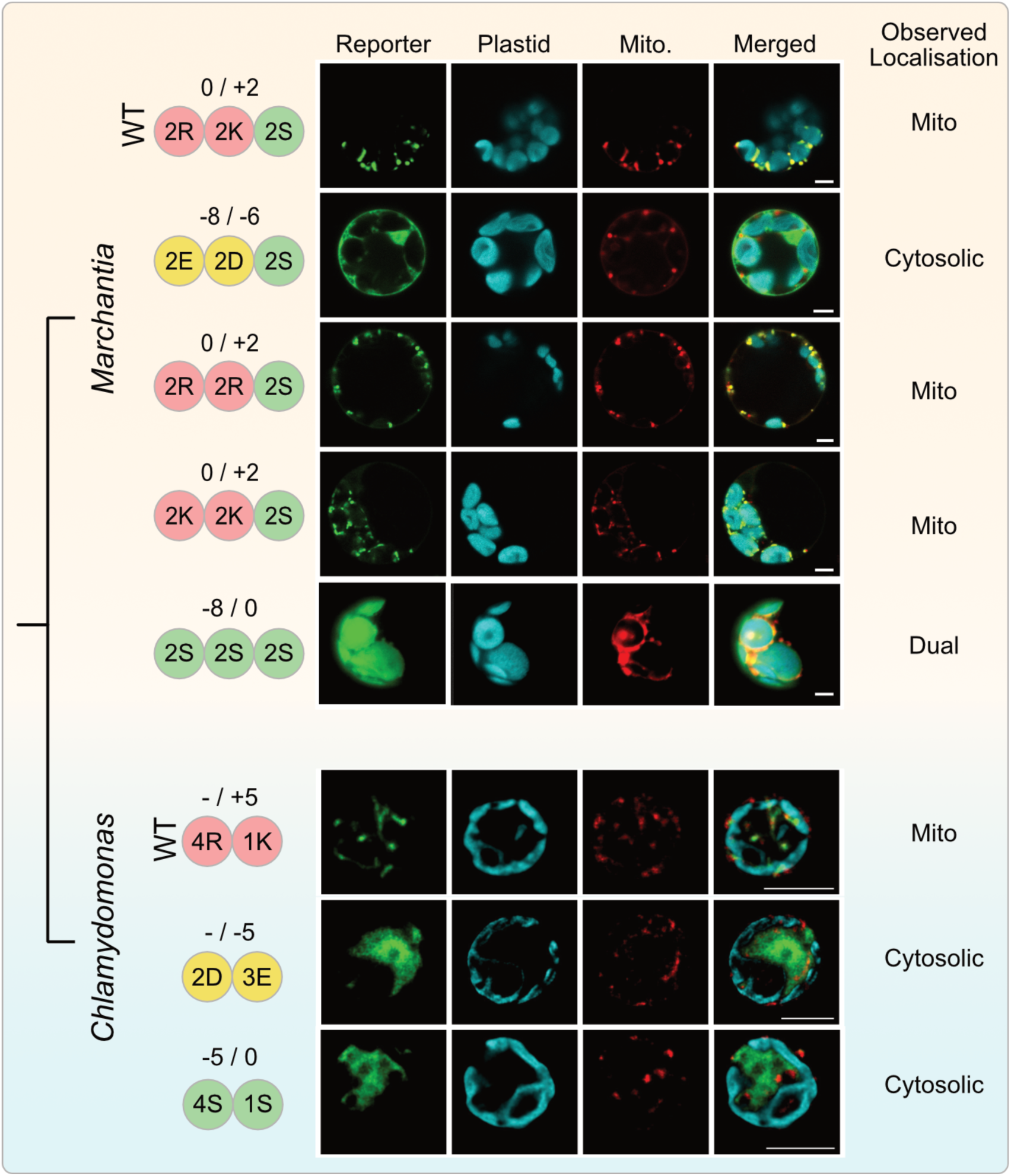
Altering only the NTS charge is sufficient to induce dual- or mistargeting. Subcellular localization of mitochondrial malic enzyme (MME1) homologues in the chlorophyte alga *Chlamydomonas* and the bryophyte plant *Marchantia*. Each row shows the localization of an independent reporter fusion and the top raw (for each species) represents the wild type (WT). Below the WT, key amino acids in the first 20 amino acid region of the NTS were substituted. The numbers of these key amino acids (in single letter codes) are shown on the left. On top of the amino acid counts, the resulting charge of the first 20 amino acids is calculated (with/out phosphorylation). Observed localization is indicated by icons on the very right. MME1 (and its variants) was localised in *Marchantia* protoplasts via citrine and *Chlamydomonas* single cells via eGFP fusion constructs. Plastids were imaged through their autofluorescence and mitochondria through MitoTracker Red. Scalebars 5µM.

Next, the idea that an amplified charged-based divergence of mitochondrial and plastid NTS was key to improving accuracy of protein targeting in land plants was tested. We used *Marchantia polymorpha*, a system to study land plant evolution^48,49^, in which the MME1 homologue (*Mp*MME1; Mp4g02270) has an overall charge of +4. As predicted, the reporter fusion with the full *Mp*MME1 sequence shows a mitochondrial localization, while the −4 mutant construct accumulates in the cytosol (Fig. 3). It is debated, whether the positive charge or amino acid identity per se per se is crucial for mitochondrial import ^50^. Substituting arginine for lysine (and *vice versa*) still leads to correct mitochondrial targeting (Fig. 3) and so for *MpMME1* a positive charge matters and not whether it is mediated by arginine or lysine. Analysis of the first 20 amino acids of 214 mNTS from 72 eudicots, where the positive charge is pronounced (above +3), shows a general preference for arginine (Fig. S6). Those proteins almost exclusively using arginine in the mNTS are bioenergetics related, while those strongly preferring lysine are mostly information processing related (Fig. S6). A preference for either lysine or arginine might hence depend on the context of protein function, which appears odd considering targeting sequences are cleaved upon import into the matrix^51^. It is feasible that arginine provides another layer of accuracy control via interaction with receptors whereas lysine only relies on charge and therefore, proteins whose targeting requires to be strictly mitochondrial might prefer arginine.

Regardless of amino acid identity, the divergence between mitochondrial and plastid NTSs (and an enrichment of serines in pNTSs) is a necessity. When the positively charged amino acids of the *MpMME1* mNTS are substituted with serine residues, the fusion construct now also targets the plastids (Fig. 3). It shows that replacing positively charged amino acids with serine residues is sufficient to dually target, or mistarget in this case, one of the key mitochondrial marker proteins. Note that a similar experiment in *Chlamydomonas* did not result in any targeting of *Cr*MME1 to the plastid (Fig. 3). This is maybe because of a requirement, of unknown nature, for a generally longer pNTS in the chlorophyte alga^52^ and a possible lack of phosphorylation in light of a late enrichment of phosphorylatable amino acids in pNTSs (Fig. 2-3).

### Origin of a membrane-anchored NTS dephosphorylase at the water-to-land transition

Two purple acid phosphatases, PAP2 and PAP9, are C-tail anchored into the outer plastid membrane in *Arabidopsis*, where they work on pNTSs^42,53,54^. *At*PAP2 (AT1G13900) and *At*PAP9 (AT2G03450)) share a 70% global sequence identity and are both characterised by the presence of an N-terminal purple acid phosphatase domain (N-PAP) and a C-tail anchor (C-TMD; Fig. 4). In addition, *At*PAP2 contains two lysine residues and a stretch of amino acids preceding the C-TMD, which, unlike PAP9, likely enables *At*PAP2 to simultaneously anchor into the mitochondrial outer membrane^54^ to also process mNTSs^55^.

**Figure 4:**
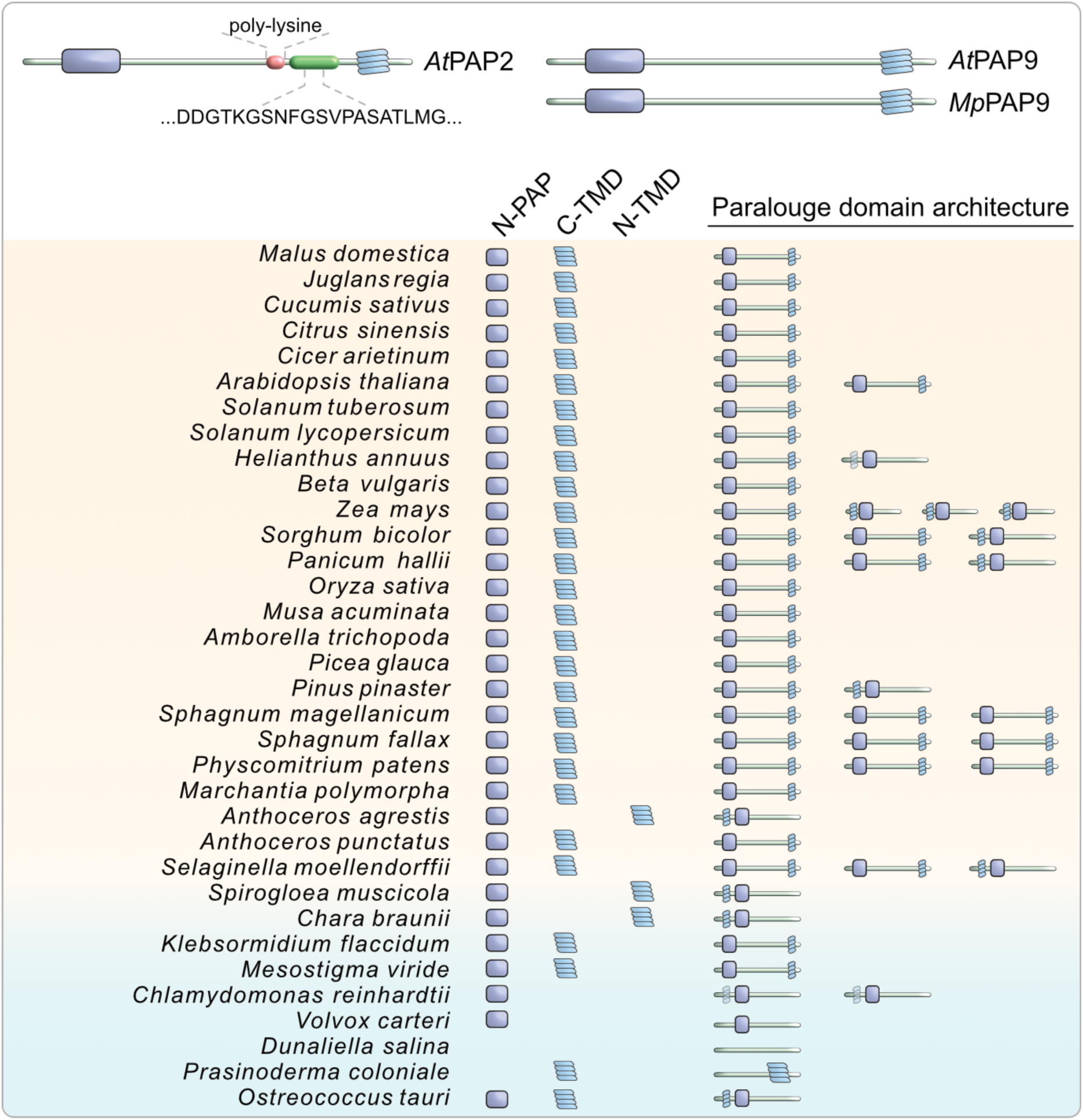
Membrane-docking of PAP2/9 evolved at terrestrialization. Domain architecture of the PAP2/9 family across various Chloroplastida. A characteristic N-terminal PAP domain (N-PAP) already evolved in chlorophyte algae, but a C-tail membrane anchor only later in streptophytes. Copy number increase in certain species is indicated by the additional icons under the column paralogue domain architecture.

While purple acid phosphatases are highly conserved across all domains of life^56^, our protein clustering identified *At*PAP2 and *At*PAP9 to be part of a smaller and likely Chloroplastida-specific family of PAPs, which we termed the PAP2/9 family. Like many other organelle related proteins, the PAP2/9 family also underwent a substantial copy number increase in embryophytes (Fig. 4). Further sequence analysis suggests that sometime during terrestrialization and commencing in streptophyte algae such as *Klebsormidium*, some of these PAP2/9 members acquired a C-TMD (Fig. 4). In some embryophytes, the members with a C-TMD duplicated and diverged further, likely giving rise to a plastid anchored PAP9 and a dually anchored PAP2.

To experimentally probe this, we focused on PAP2/9 homologues of *Chlamydomonas* and *Marchantia* (Fig. S7). A candidate *Chlamydomonas* PAP2 homologue (Cre11.g468500) has a predicted transmembrane domain only at the very N-terminus (ca. 1-30 amino acids), which is likely a part of a targeting sequence (Fig. S7). N-terminal membrane docking of proteins is rare, while N-terminal targeting sequences are the norm^57^. Reporter fusion of this N-terminal region alone led to a mitochondrial (likely matrix) localisation of *CrPAP2* (Fig. S7), hinting at the ancestral localisation of the PAP2/9 protein family represented by chlorophyte algae. *Marchantia* encodes only a single C-TMD containing PAP (Mapoly0122s0035), which lacks the characteristic amino acid stretch present in *AtPAP2* but encodes a C-TMD (Fig. S7). We therefore refer to this protein as *Mp*PAP9. *Mp*PAP9 localizes to the plastid (Fig. S7) and likely captures an intermediate state of PAP2/PAP9 protein family evolution, where the acquisition of a C-TMD lead to the origin of a plastid outer membrane anchored PAP.

### Remodeling of import platforms alongside plant diversification

To trace the evolutionary history of import components across the Chloroplastida, we clustered whole proteome models of 42 species (Table S4C) based on sequence similarity ^58^ and screened the clusters (Table S4B) for known import components (Table (S4C-D). The result shows that many components of mitochondrial and plastid protein import were duplicated around the time of plant terrestrialization (Table S4E, Fig. S8), i.e. from the zygnematophyceaen algae onwards. This is in line with a more general pattern regarding the expansion of mitochondrial and plastidal functions during the transition from water to land^23^.

At the inner mitochondrial membrane, duplication and divergence of TIM23 (Fig. 5A) led to isoforms such as TIM23-II, which in *Arabidopsis* allows the finetuning of protein import in response to changing levels of respiratory complex I occurring e.g. under oxidative stress ^59^. TIM21 was also duplicated, with the copies likely also interacting with proteins of the electron transport chain ^60^. Peripheral components of TIM23, TIM22 and the TIM23-17 complex at large, such as PRAT (Preprotein and Amino acid transporters) also duplicated in the green lineage (Fig. 5A, Fig. S8). These newly recruited components, which connect protein import with respiration and oxidative stress, likely benefited a life on land where oxidative stress is more pronounced. At the outer mitochondrial membrane, the receptors TOM20 and TOM22, and the main import channel TOM40, also significantly diverged from their non-photosynthetic homologues^61^. One reason could be the need for better cargo selection now that the plastid endosymbiont was present with its own emerging import apparatus, namely TOC/TIC^43^. That is in accordance with the *Arabidopsis* mitochondrial NTSs (mNTS), unlike those of animals, displaying two hydrophobic domains that bind to two separate binding sites of the *At*TOM20 ^62–67^. In combination, the changes at the two mitochondrial membranes likely improved import rate, accuracy and regulation.

**Figure 5:**
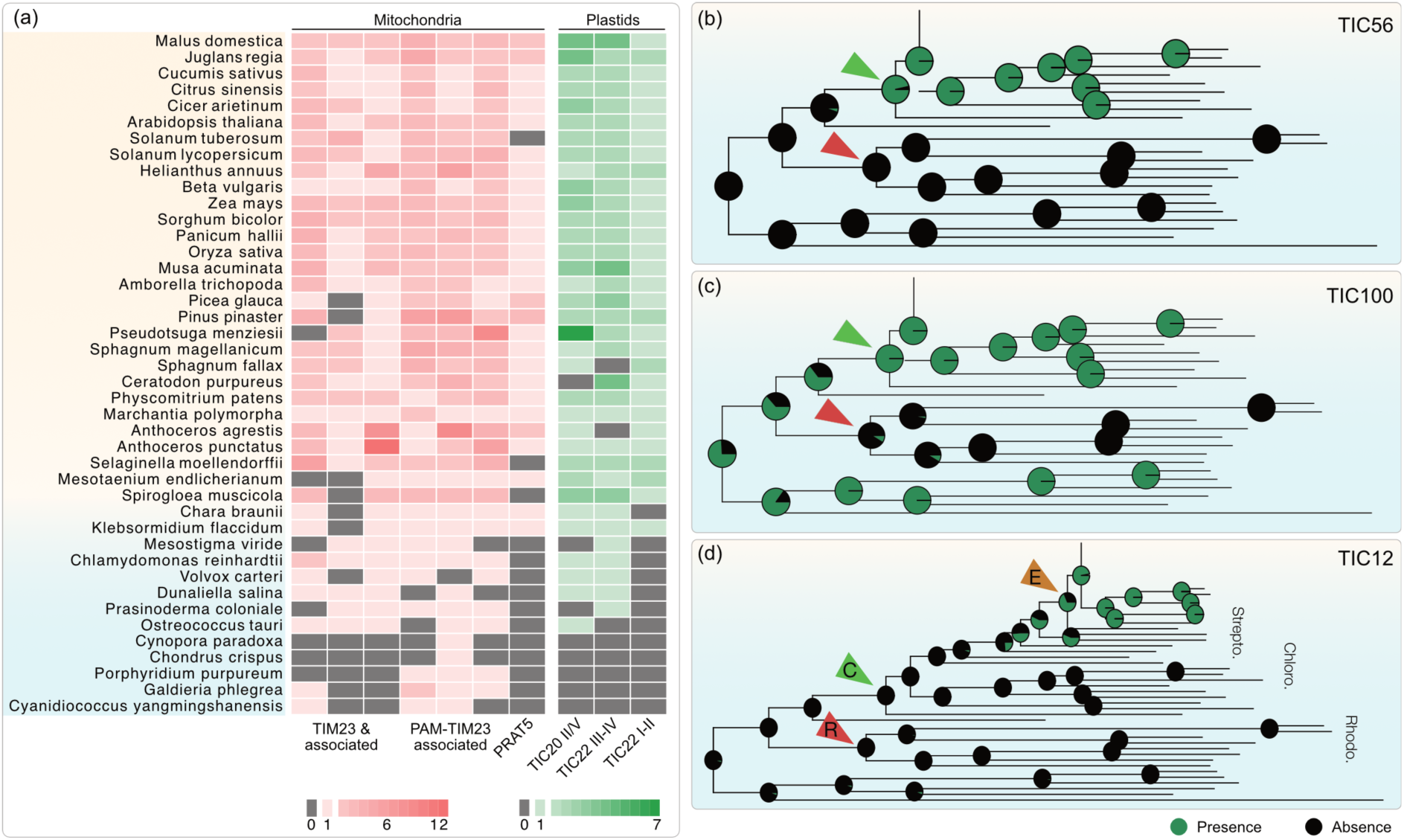
Recruitment, expansion and divergence of import receptors. **(a)** Gene copy numbers of key import channels at the inner membranes of mitochondria (crimson) and plastids (green) (n=42 Archaeplastida species). **(b)** Ancestral state reconstruction for the key components of TIC complex (n=137 Archaeplastida species, 6 outgroups). Ancestors of rhodophytes, chlorophytes and embryophytes are shown in red, green and mustard. The phylogeny of the speices used and full ASR of the three import components is shown in Fig. S9-12.

At the inner plastid membrane, duplications of TIC20, TIC62 and TIC22 (Fig. 5) gave rise to quantitative and qualitative differences in plastid import. For instance, a duplication of TIC22 provided functionally redundant isoforms that increase the rate of import in land plants ^68^. In contrast, isoforms resulting from duplications of TIC20 contributed to a qualitatively different import complex, dedicated to import of photosynthetic proteins, termed “photosynthetic-type” TIC complex (characterised by TIC20, TIC56 and TIC214)^69^. While TIC20-IV formed the core of this complex, a massive divergence of another TIC20 isoform likely resulted in TIC214 with only a shared membrane-spanning region between TIC20 and TIC214 ^70^. After its emergence in Chloroplastida, as also substantiated by its presence in *Chlamydomonas* ^71,72^, this complex was further modified in land plants via addition of TIC12 (Fig. 5D), a recently identified component of this complex^73^.

On the whole, our bioinformatic analyses and experimental validation suggest that ever since an algal’s transition from water to land, coevolution of receptor platforms (particularly at the inner membranes) and NTS, as well as divergence of mNTS and pNTS, resulted in improved accuracy (and likely also rate) of protein import by the two key organelles of plants.

## Discussion

The eukaryotic cell is required to orchestrate the correct targeting of thousands of cytosolically translated proteins. Eukaryotes, especially photosynthetic ones, have therefore evolved a variety of mechanisms that contribute to the correct targeting and import selection: organellar mRNA localisation^74,75^, alternative transcription and translation initiation^76^, ubiquitination^77^, piggy-back transport^78^, phosphorylation^43,79,80^ and a plethora of small GTPases when taking vesicle trafficking as a mean to specifically transport proteins from one destination to another into account^81,82^. Some are not regulated but work constitutively, while others secure protein homeostasis of a compartment or to swiftly respond to environmental changes^83^. Each import strategy adapts according to the requirement of a specific evolutionary niche and conversely, changes in protein targeting can offer novel possibilities.

The expansion of plastid proteomes underpinned the evolutionary success of the chloroplastidal lineage and its subsequent adaptation to the land and its novel stressors^23,84^. Concomitant changes of the import machinery, such as a duplication of the main outer membrane import channel TOC75, allowed for a better rate of import of an increasing amount of cargo, which in return, is likely associated for instance with a better response to high light stress in the green lineage^47^. Similarly, the expansions of plastidal and mitochondrial inner membrane channel components (Fig. 6, Fig. S8) likely increased import rate, which can help to adapt to altering levels of oxidative stress by better regulating import.

**Fig. 6:**
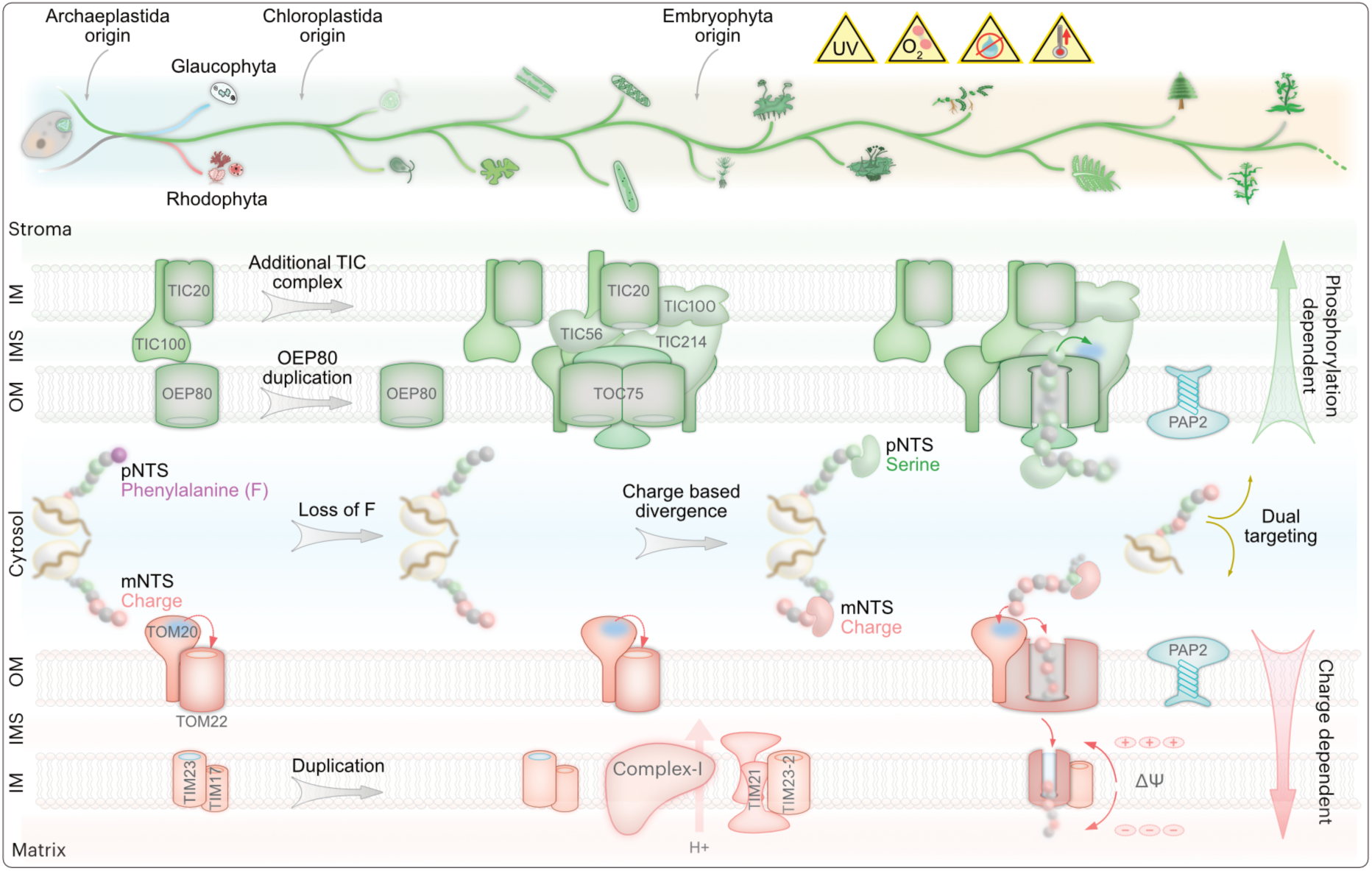
Schematic of mitochondrial and plastid protein import evolution. For simplicity, a linear evolution of the main archaeplastidal lineages is pictured at the top. Chloroplastidal diversification on land was characterised by organellar adaptation towards various stressors^23^ and a recruitment of novel proteins that require import. At the origin of the Archaeplastida, a conserved phenylalanine motif in the pNTS evolved for plastid targeting. It was lost at the origin of the green (chloroplastidal) lineage, arguably also leading to protein mistargeting between plastids and mitochondria ^47^. This, along with the expansion of the organelle proteomes, particularly at the water to land interface ^23^, imposed a selection pressure for the remodeling of import machinery towards improved rate, accuracy and regulation. Duplication and divergence of key import channels (e.g. TOC75, TIC20, TIM23), giving rise to components such as an additional TIC complex, allowed the import of expanded organelle proteome, likely at an increased rate, facilitating adaptation to light and oxidative stress^47^. On the cargo side, the first 20 amino acids of N-terminal targeting sequences for plastids (pNTS) and mitochondria (mNTS) diverged, whereby pNTS accumulated phosphorylatable amino acids (depicted in green) and averted positively charged amino acids (depicted in salmon). As a consequence, plastids and mitochondria could more accurately differentiate and import their cargo, in phosphorylation and charge dependent manner. The evolutionary divergence of NTSs was undergirded by Δψ and a coevolution with receptors by mechanisms such as (i) initial, and likely ancient, charge-based screening by the outer membrane receptor TOM20/44, and (ii) Δψ-dependent activation of a cation selective channel TIM23, screening for positively charged mNTSs on the matrix side. Chaperons can also guide cargo to organelles, likely improving accuracy^80,89–91^. Membrane recruitment of a dephosphorylase such as PAP2 further contributed to the regulation of (dual) targeting.

Co-evolution of import receptors and N-terminal targeting sequences (NTS) also contributed to improved cargo selection. For instance, TOM20 and TOM22, which mediate the entry of the mNTS ^85^, appear to be substantially diverged in Archaeplastida and mechanism of cargo selection differ between plants and animals. In animals, the cytosolic domain of TOM22 interacts with the mNTS and imports it further. The plant TOM22 significantly differs and instead serves as a gate-keeper by occupying a hydrophobic cradle on TOM20 ^85–87^. TOM22 gets outcompeted by mNTSs and displaced – but likely not by a pNTS – to allow mitochondrial cargo entry through TOM20 ^85–87^. TOM22 interacts with the mNTS via a negatively charged region and likely selects positively charged mNTSs in the glaucophyte alga *Cyanidioschyzon*^45^, once more underscoring the relevance of charge-based selection for organelle cargo at the outer mitochondrial membrane in early-branching phototrophic eukaryotes. At the inner mitochondrial membrane, TIM23 forms a cation selective channel that is activated by the mNTS and membrane potential (Δψ) of the inner mitochondrial membrane^88^. And finally, Δψ can, more directly, facilitate entry of positively charged mNTSs, while averting the negatively charged NTS such as a phosphorylated pNTS of land plants^43^. At least two major receptors at the inner and outer membranes, along with Δψ, hence contribute to a selection pressure that supressed positive charges in pNTSs. It is this divergence that our substitutions erase, resulting in the dual targeting of a mitochondrial marker protein (Fig. 3).

On the plastid side, changes at the inner envelope membrane and in the pNTS also improved accuracy protein import. It is feasible that the chloroplastidal recruitment of the TIC214-containing import complex (Fig. 5) re-installed the import accuracy that was lost with the loss of a phenylalanine-based NTS selection at the origin of the Chloroplastida^47^. Molecular dynamics simulation on the TIC-TOC super complex structure predicted an interaction of the pNTS with TIC214 ^72^, which was confirmed by cross-linking experiments that revealed an interaction of the rbcS NTS with positively charged residues of TIC214 in its preprotein translocation path ^71^. Such an interaction is likely to repeal the generally positively charged mNTS and select a pNTS for further import of the protein.

Evolving many phosphorylatable sites in a pNTS only serves a purpose when accompanied by appropriate kinases and phosphatases. The kinase STY17 in *Arabidopsis* phosphorylates pNTSs, which is then passed onto OEP61 via HSP70-14/3/3 guidance complex and imported at an increased rate ^39–41,80,89,92,93^. The evolutionary history of such chaperons, kinase and phosphatase remains blurry, owing to their generally conserved nature and abundance. By combining protein clustering, domain analyses and experimental approaches, however, we were able to verify the embryophytic recruitment of at least one outer plastid membrane-localised dephosphorylase, a member of the PAP2/9 family, which plays a role in protein import in *Arabidopsis* ^42,55^. Along with a concomitant enrichment of phosphorylatable amino acids in the pNTS, particularly pronounced in angiosperms (Fig. 1-2), a recruitment of a plastid outer membrane dephosphorylase (Fig. 4) hints at a layer of differential targeting control to mitochondria and plastids still poorly understood (Fig. 6).

A duplication of this membrane-anchored phosphatase, likely in angiosperms, gave rise to a paralog such as *At*PAP2, which is dually localised to plastid and mitochondrial outer membranes. Since phosphorylated amino acids in the pNTS is thought to act as a mitochondria aversion signal^43,94^, the presence of a dephosphorylase on the mitochondrial outer membrane appears to functionally reverse the evolutionary changes, potentially allowing for organelle mis- or dual-targeting. Indeed, a known substrate of *At*PAP2 is a dual targeted protein, mitochondrial import of which hinges upon de/phosphorylation ^55^. Some 110 proteins in *Arabidopsis* are known to be dually targeted ^95,96^ and the dNTS physicochemical features represent an intermediate value between those of pNTSs and mNTSs^43^.

The relative contributions of NTS secondary structure and NTS physico chemical features to correct targeting are a matter of ongoing investigations, especially due to a lack of distinct features other than the abundance of serines in pNTS and alpha helices in mNTS^91,97–100^. The MME1 (a mitochondrial enzyme) wildtype sequences (*Cr*MME1 and *Mp*MME1) used in this study are predicted to have an alpha helix in the first 20 amino acids (Fig. S13). However, the *Marchantia* MME1 with six serines (with an intact alpha helix) indeed localises to the plastid and provides an example, where physico-chemical features surpass the secondary structure to convert a mitochondrial protein into a dual targeted protein. To what extent this could be a way towards dual targeting remains to be addressed along with the role of de/phosphorylation (by, e.g. PAP2), and the mechanism for the membrane recruitment of dephosphorylases (e.g. a direct docking or via the interaction with import receptors/cytosolic factors).

We propose that an early ‘mis-sorting’ due to a loss of phenylalanine at the origin of Chloroplastida, later partly evolved into (regulated) protein dual targeting via a coevolution of NTSs, import receptors and components such as PAP2/9. Experimental information regarding protein dual targeting outside of angiosperms, however, is rare and the prediction of dual targeted proteins remains unreliable ^101^. Studying the origin and evolution of (regulated) protein dual targeting is hence a promising avenue and calls for more dedicated research outside of angiosperms.

In closing, our computational analyses across a wide range of photosynthetic eukaryotes underscores major shifts in the NTSs from algae to angiosperms. The sample size of reliable genomes and organelle proteomes is skewed towards embryophytes and affects our inference as to where along the evolution of plants the NTS diverged exactly. For instance, it is conceivable that NTSs started to diverge immediately after the loss of phenylalanine at the origin of the Chloroplastida^47^. A better resolution was not achieved, owing to a combination of poor sample size and the likely strong selection pressure in evolving novel means of separating mitochondrial from plastid cargo. Additional genomes, organelle proteomes, and experimental studies on protein import in algae (including rhodo– and glaucophytes), will aid in tracing the evolution of organelle NTSs at a finer resolution. Notwithstanding the current limitations, it is clear that the divergence of physico-chemical properties of NTSs played a crucial role in preventing protein mistargeting (Fig. 2-4) and it co-evolved with their processing counterparts at the outer membrane (Fig. 5). In light of the loss of phenylalanine from the pNTS ^47^ and a general lack of conserved determinants between the two NTSs^91,99^, this divergence allowed for an improved recognition of cargo at the outer membranes of organelles of endosymbiotic origin (Fig. 6) and facilitated an accurate import of the expanding organelle proteomes that coincided with the diversification of plants^23^.

## Conclusion

Contributions of mitochondria and plastids to plant’s adaptation and divergence are plenty ^23,84,102^. In this study, we focused on the evolution of the organelle protein import itself, a gatekeeper for the organellar contributions to a species fitness. Origin and diversification of the Chloroplastida, particularly that of embryophytes, was facilitated by major changes in import, also evident in targeting sequence divergence between mitochondrial and plastid cargo. Experimentally contrasting organelle protein import in the single cell alga *Chlamydomonas reinhardtii* with that in the early diverging bryophyte *Marchantia polymorpha*, we validate the importance of targeting sequence divergence with respect to charge for enhancing targeting accuracy. Our revised model highlights novel mechanisms that determine import fidelity and regulation, such as de/phosphorylation at the outer membranes and ΔΨ at the inner mitochondrial membrane; mechanisms not only confined to plants, but also expandable to mitochondrial-nuclear localisation shifts in humans where phosphorylation of a transcription factor was recently shown to be responsible for maternal inheritance of mitochondrial DNA^79^. Membranes, ΔΨ and phosphorylation are traits likely as ancient as life itself. How their interplay has evolved into a selection and regulation mechanism for protein cargo transfer across membranes of endosymbiotic origin presents a promising avenue of future research.

## Materials and methods

### Genomes used in the study

To study the evolution of NTS and import components, an inhouse database of 137 Archaeplastida and 6 sister species was consolidated from available databases (species names and sources summarised in Table S1A). NTS analyses were conducted on 102 genomes from major clades where organelle proteomes could be reliably inferred based on available experimental proteomes (Table S1B). The presence-absence patterns of import components were plotted for a subset of genomes (Table S1C), representative from each clade, to visualise the broad patterns clearly. The ancestral state reconstructions (ASR) of import components were conducted on all genomes.

### Analyses of NTS from algae to angiosperms

To study the features of NTS across organelle targeted proteins, we used 102 Chloroplastida species from five major clades: Eudicots (66), Monocots (14), Bryophytes (7), Streptophyte (6) and Chlorophytes (9) algae (Table S1B). Within-clade all v. all reciprocal best blasts hits (rbbh) were retrieved for all proteins from all the species of these clades. To infer organelle proteomes of all species in a given clade, we used experimental plastid and mitochondrial protein (Gene IDs of the experimental proteins summarised in Table S2A-C) available across Chloroplastida^103–105^. For a protein in a given species, if a rbbh was found against an experimentally validated organelle protein in the same clade, we inferred that protein to be organelle localised for that species (Gene IDs of inferred proteins summarised in Table S3, method schematic in Fig. S1). As there are no experimental proteomes available for streptophyte algae, we inferred their organelle proteomes based on chlorophyte algae. From these inferred plastid, mitochondrial and dual targeted proteomes, we analysed the first 20 amino acids for electric charge and content of phosphorylatable amino acids of each protein using (the script, the resulting 20 AA sequences across species, charge/phospho values of each protein across all species are available on Zendeo; means for all species are consolidated in Source Data Fig. 1 and plotted in Fig. 1).

To investigate ancestral states of these two traits, a phylogenetic tree of each clade was constructed using concatenation of organelle proteins present in all species in that clade (sequences and Newick files available on Zenedo), in IQ-TREE^106^ v2.0.3 after aligning them with MAFFT^107^ v7.505. The resulting unrooted trees were rooted using minimal ancestral deviation ^108^. Using this phylogenetic tree for the clade and based on values of charge and phosphorylatable amino acids across all species in the clade, ancestral state reconstruction (ASR) was conducted in rstudio (Phytool^109^ package 0.7.80) (individual outcomes summarised on Zenedo; means consolidated in Source Data Fig. 2 and plotted in Fig. 2 as well as in Fig. S3 and Source Data Fig. S3).

### Analyses of PAP2/9 homologues

PAP2/9 homologues from all species were analyzed through Interproscan^110^ and DeepTMHMM^111^ to annotate functional domains and membrane anchoring domains, respectively.

### Cloning and transfection

Genes of interest from *Marchantia, MpMME1* (Mp4g02270) and *MpPAP9* (Mapoly0122s0035) were amplified from the cDNA, were cloned into *Gateway* (Invitrogen) entry vector pDONR221^TM^ (Invitrogen). Substitutions of interest were made into the wild type MME1 using PCR-mutagenesis. All constructs were transferred at the N terminal of Citrine on pMpGWB106^112^, or at the C terminal of citrine on pMpGB105 (for *MpPAP9*) via *Gateway* cloning. The N terminal of interest from *Chlamydomonas CrMME1* (Cre06.g268750) and *CrPAP2* (Cre11.g468500), were invitro synthesized and amplified from the genomic DNA, respectively. Subsequently, they were cloned at the N terminal of eGFP into the vector pJR38^113^ via ndeI-bglII (bglII site was introduced to pJR38 prior to cloning the sequences of interest).

Localization vectors were integrated into *Marchantia* genome via *Agrobacterium*^114^. Briefly, the vectors were first introduced into *Agrobacterium tumefaciens* (GV3101 without pSOUP) by electroporation (Bio-Rad GenePulser Xcell, 1.44 kV). Epical notches were cut away from two-week-old thalli and after two days of recovery, these explants were incubated with *Agrobacterium* (starting OD_600_ 0.02) for three days. After washing several times with water, the explants were plated onto Hygromycin (10 μg/mL) and Cefotaxime (100 μg/mL) where the explants were selected and further propagated. The constructs were introduced to *Chlamydomonas* via the glass beads method^115^. Briefly, 1µg of linearized plasmid, 5% PEG6000, 330µL *Chlamydomonas* culture (concentrated to 10^8^ cells per ml) and 300mg glass beads were vortexed for 15 seconds. The cells were plated, selected and further propagated on 10µg/mL Paromomycin. The algal and plant lines used in the study are summarized in Table S5.

### Plant and algal growth conditions

*Marchantia polymorpha* ecotype Takaragaire-1 (Tak-1) were grown on half-strength Gamborg B5 agar-plates under continuous light (70 μmol m^−2^ s^−1^). *Chlamydomonas reinhardtii UVM4* ^113^ was grown in high salt medium supplemented with acetate as a carbon source and were grown under continuous light (40 μmol m^−2^ s^−1^).

### Protoplast isolation

300-500mg of two-week-old *Marchantia* thalli were incubated in 8% Mannitol, 6.3 g/L Gamborg B5 Vitamins (MGV) with 15mg/ml Cellulase and 5 mg/mL Macroenzyme R10 overnight. Protoplasts were filtered with 50µm strainer and washed twice with MGV by centrifuging at 300g, 3min and gently resuspending by swirling the tube.

### Microscopy

*Marchantia* protoplasts and *Chlamydomonas* single cells were strained with 300nM MitoTracker™ Red CMXRos as per manufacturer’s instructions to visualize mitochondria. Plastids were visualized via chlorophyl autofluorescence at 480/590 nm. GFP/citrine reporters were visualized at 470/520 nm.

### Origin and evolution of import components across Archaeplastida

42 Archaeplastida species (Table S4A), from each available representative clade, were clustered using OrthoFinder version 2.5.4 ^58^, after all vs all blasts were conducted (E-value cutoff 10e-6) using diamond blast version2.011^116^. The protein clusters (Table S4B) were annotated based on the presence of experimentally validated import components (Table S4C-D), across species gene copy numbers (GCN) for each component were inferred (Table S4E) and a subset was plotted in Fig. 5A (see Fig. S8 for GCN plots of all components).

To investigate the origins of a select components more rigorously, we utilised the full database of 137 Archaeaplastida species with 6 outgroup species (Table S1A). Considering the sequence divergence of these components, iterative hidden Markov searches were conducted using HMMER^117^ 3.3.2 and seed sequences from Arabidopsis. The homologues thus retrieved were used for ASR in rstudio (Phytool^109^ package 0.7.80).

## Data availability

Supplementary figures and tables are available with this submission. Additional data are available on Zenedo (https://zenodo.org/uploads/14988946) which includes, but is not limited to, the following supplementary data and in-house scripts:

Source Data Figure 1-5

Source Data Figure S2-13

Table S1 Species name, taxonomy and source of genomes used in the study

Table S2 Gene IDs of experimental proteomes used as seeds to infer organelle proteomes

Table S3 Gene IDs of inferred organelle proteomes

Table S4 Gene IDs of experimentally validated import components and their inferred gene copy number rbbh.py Conduct all vs all BLAST for each clade

Organelle_proteome_inference.py Infer organelle proteomes based on rhhb results and experimental proteomes

getNTS1-20AA.py Get the first 20 amino acids of inferred organelle proteins

getNTS1-20AA_charge_phospho.py Get charge and phosphorylatable amino acids from the first 20 amino acids of inferred organelle proteins

NTS_charge_phospho_plots_stats.py Summaries and plot averages of charge and phosphorylatable amino acid content across species and clades

clade_phylogeny.py Infer phylogenetic trees for all clades

charge_phospho_for_ASR.py Prepare the charge and phosphorylatable amino acid content data for ASR

NTS_ASR.r Infer ASR for charge and phosphorylatable amino acid content of each clade

NTS_ASR.r Infer ASR for import components of interest

## Author contributions

PKR: Conceptualization, Supervision, Experimental design, Methodology, Investigation, Data curation, Validation, Formal analysis, Visualization, Writing - original draft, review and editing. CGG: Investigation, Data curation, Validation MDSS: Investigation, Data curation SBG: Conceptualization, Project administration, Supervision, Experimental design, Funding acquisition, Resources, Visualization, Writing - original draft, review and editing.

## Funding

We thank the Deutsche Forschungsgemeinschaft (SFB 1208-2672 05415 and SPP2237–440043394) for funding to SBG.

## Acknowledgments

We acknowledge support from the high-performance computing cluster (HILBERT) at the Heinrich Heine University Düsseldorf and thank Michael Knopp, Paula Tief, David Mohr, Kira Tormo Arlt and Margarete Stracke for their help. We also thank Prof. Hirofumi Nakagami and Katharina Kramer of the Max Planck Institute for Plant Breeding Research (Köln, Germany) for guiding our experiments in *Marchantia polymorpha*. CGG and PKR are grateful to Prof. William F. Martin for providing a financial support.

## Notes

https://zenodo.org/uploads/14988946

### Competing Interest Statement

The authors have declared no competing interest.

