## Supplementary Figures 1-13 for "Evolutionary refinement of mitochondrial and plastid targeting sequences coincides with the late diversification of land plants"

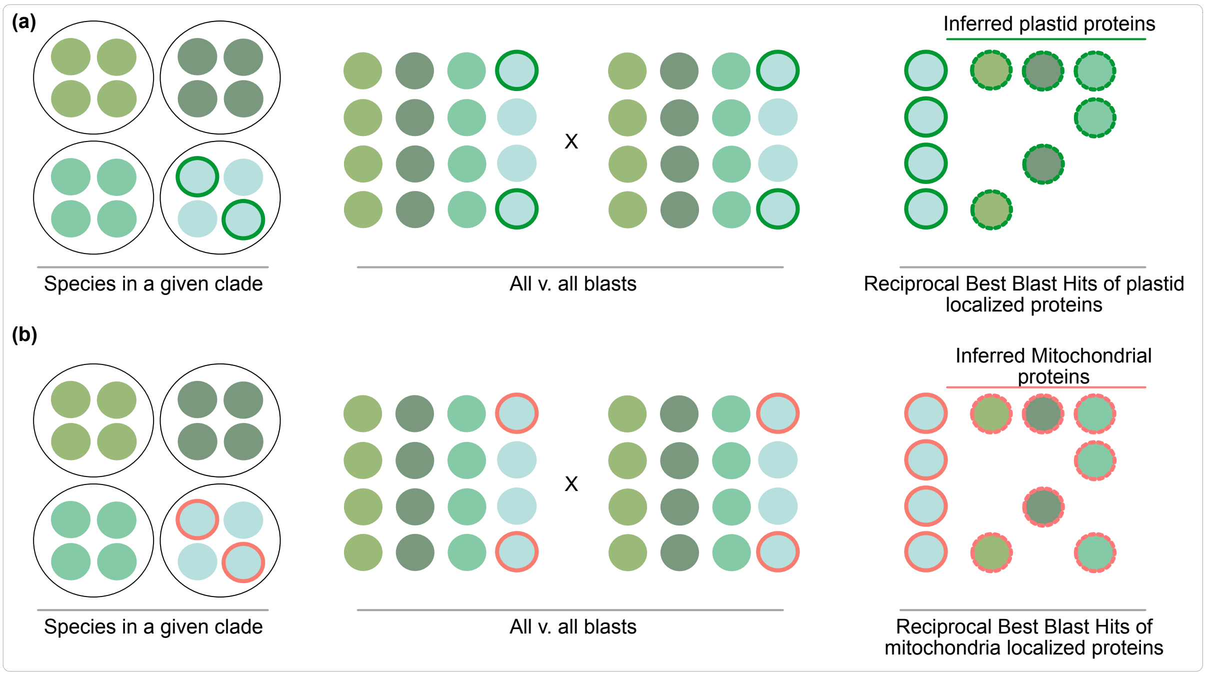


**Fig. S1: Inferring organelle localised proteins across five major chloroplastida clades.** 110 Archaeplastida species of chlorophyte and streptophyte algae, bryophytes, monocots and eudicot clades were used for the NTS analysis. Each black circle outline represents a species and the smaller, coloured dots represent individual proteins. Dots with coloured outlines represent experimentally validated plastid (green outline) and mitochondrial (salmon outline) proteins. All vs. all blasts were conducted for all proteins, from all species within each clade. Reciprocal best Blast hits (RBBH) were obtained for all experimentally validated **(a)** plastid- and **(b)** mitochondria-localised proteins in a given species to infer organelle localised proteins (dotted outlines, on the right) for all species in that clade.

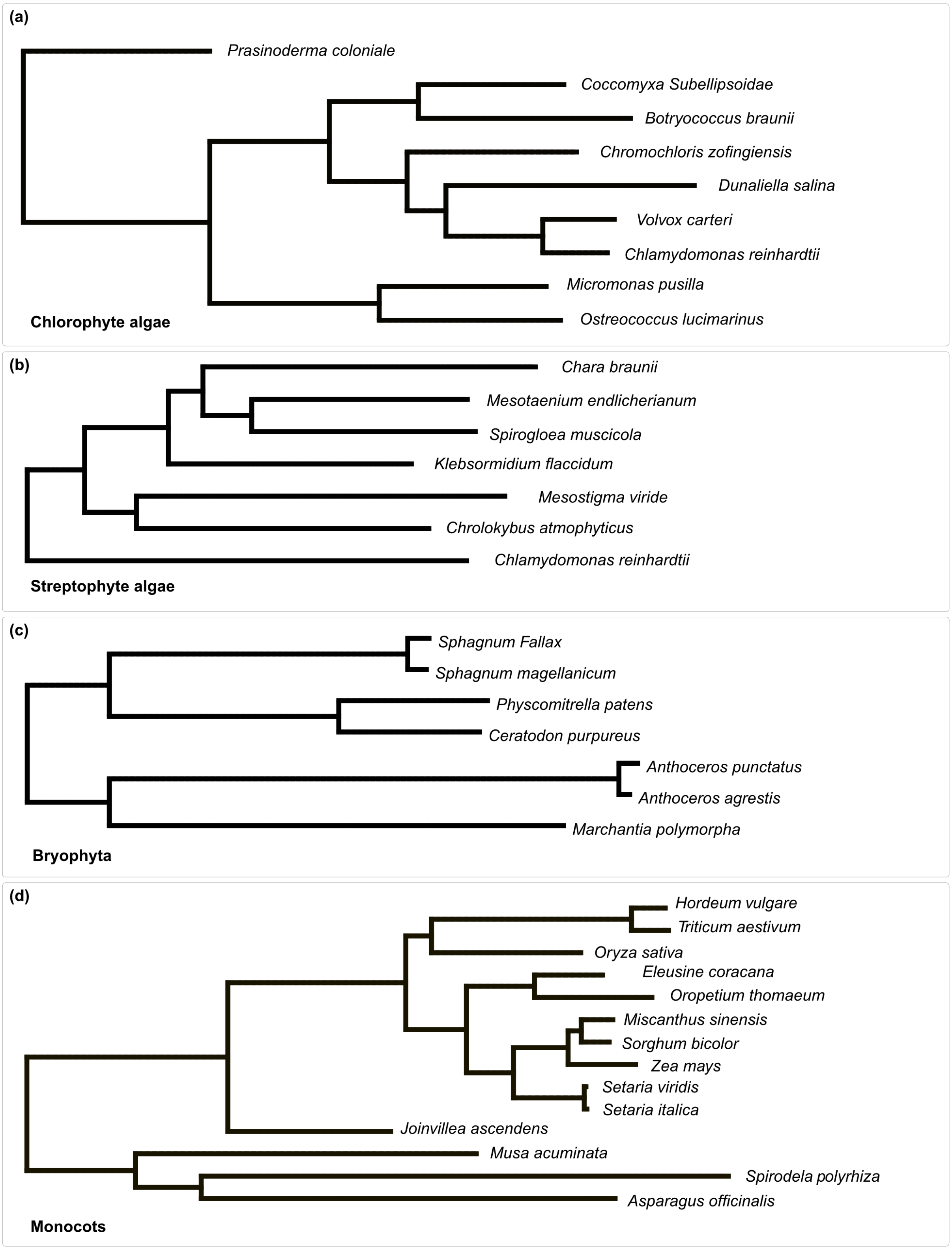
(a)
Prasinoderma coloniale
Coccomyxa Subellipsoidae
Botryococcus braunii
Chromochloris zofingiensis
Dunaliella salina
Volvox carteri
Chlamydomonas reinhardtii
Micromonas pusilla
Ostreococcus lucimarinus
Chlorophyte algae
(b)
Chara braunii
Mesotaenium endlicherianum
Spirogloea muscicola
Klebsormidium flaccidum
Mesostigma viride
Chrolokybus atmophyticus
Chlamydomonas reinhardtii
Streptophyte algae
(c)
Sphagnum Fallax
Sphagnum magellanicum
Physcomitrella patens
Ceratodon purpureus
Anthoceros punctatus
Anthoceros agrestis
Marchantia polymorpha
Bryophyta
(d)
Hordeum vulgare
Triticum aestivum
Oryza sativa
Eleusine coracana
Oropetium thomaeum
Miscanthus sinensis
Sorghum bicolor
Zea mays
Setaria viridis
Setaria italica
Joinvillea ascendens
Musa acuminata
Spirodela polyrhiza
Asparagus officinalis
Monocots

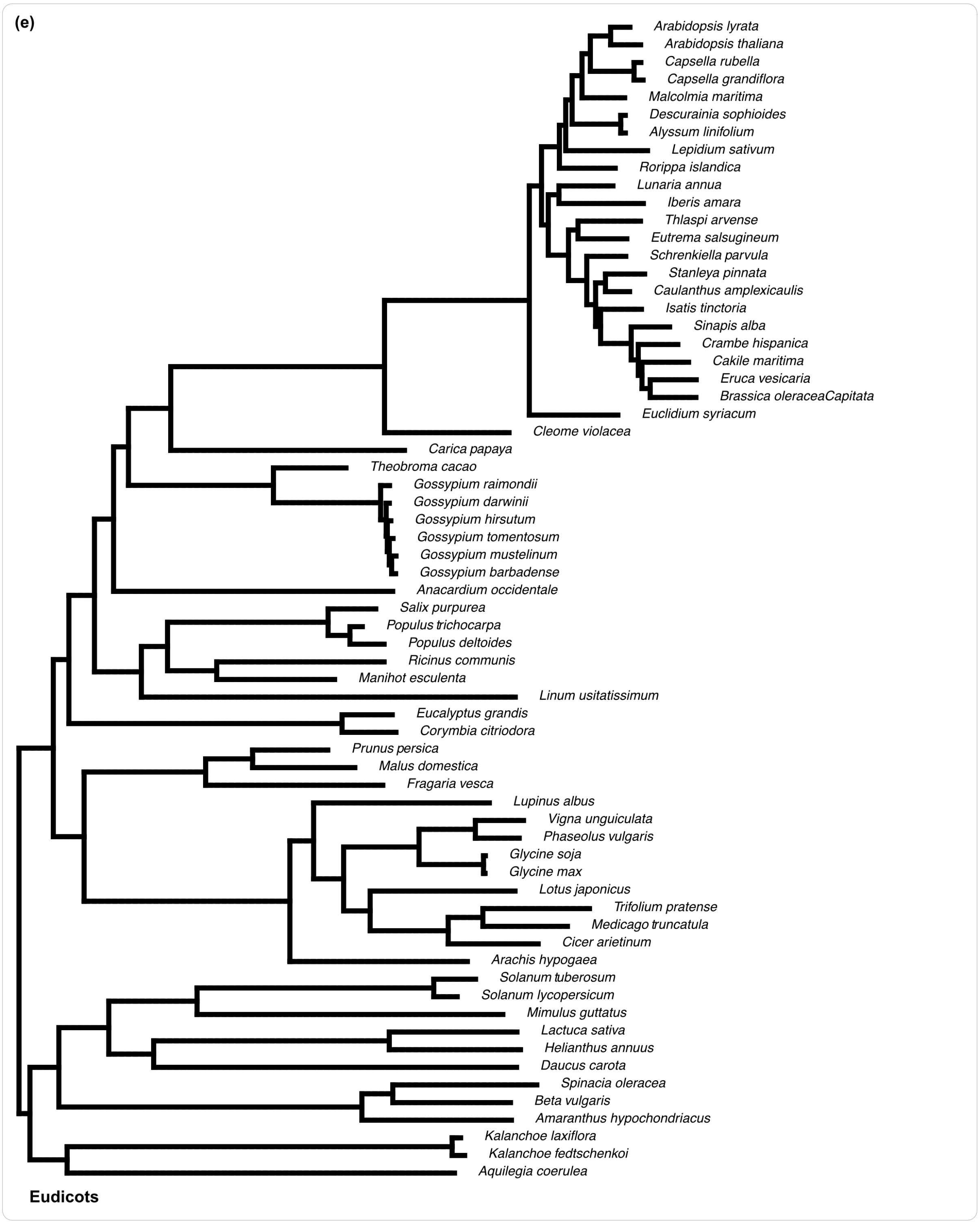


**Fig. S2: Phylogenies of the five major chloroplastidal clades**. Inferred phylogenies of each clade based on the concatenation of organelle localised proteins present in all species for that clade.

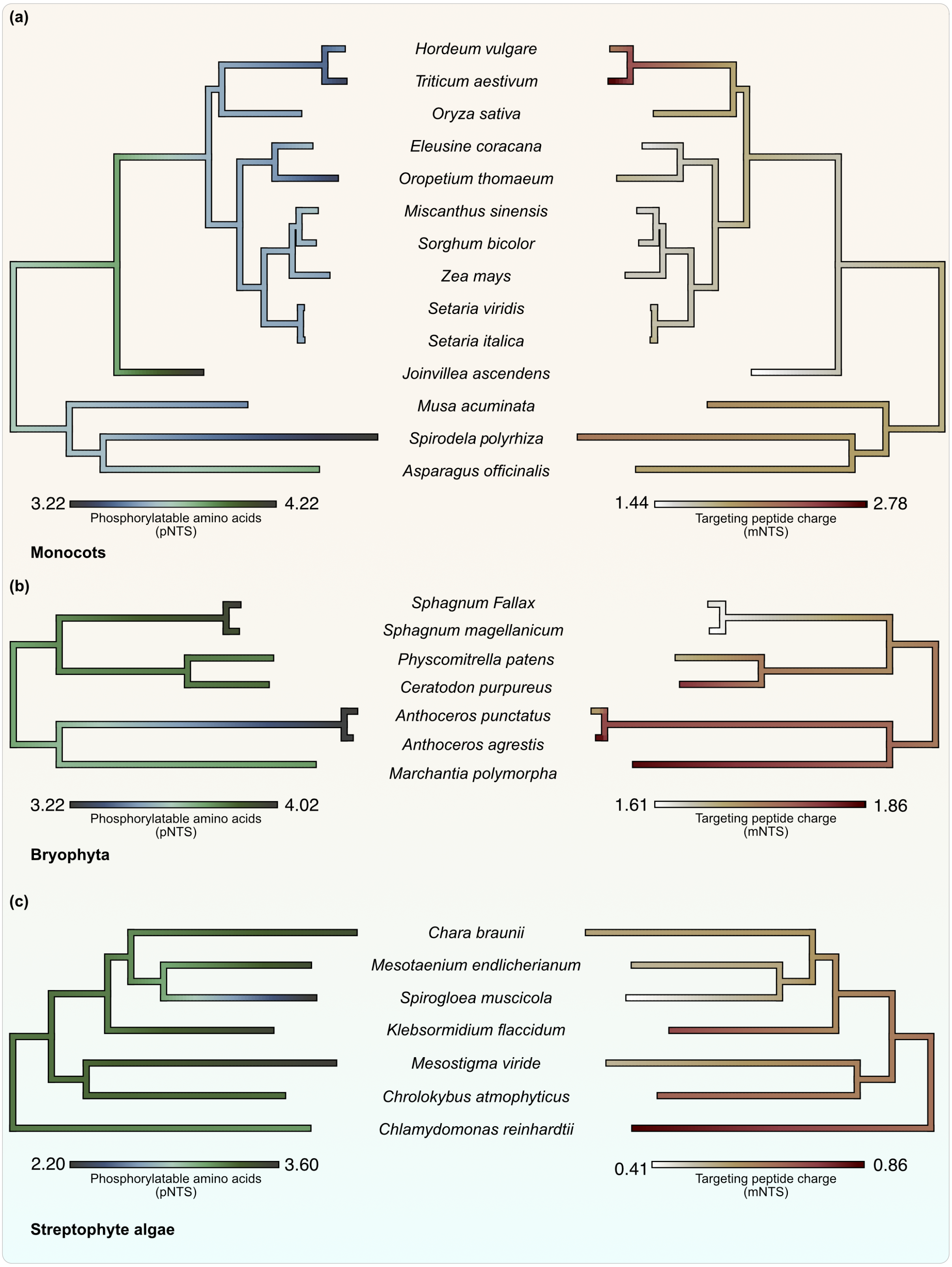


**Fig. S3:** Ancestral states of phosphorylatable amino acids in pNTSs and charges of mNTSs, inferred across ancestors of **(a)** monocots, **(b)** bryophytes and **(c)** streptophyte algae; color-coded based on the values of the two features across ancestors.

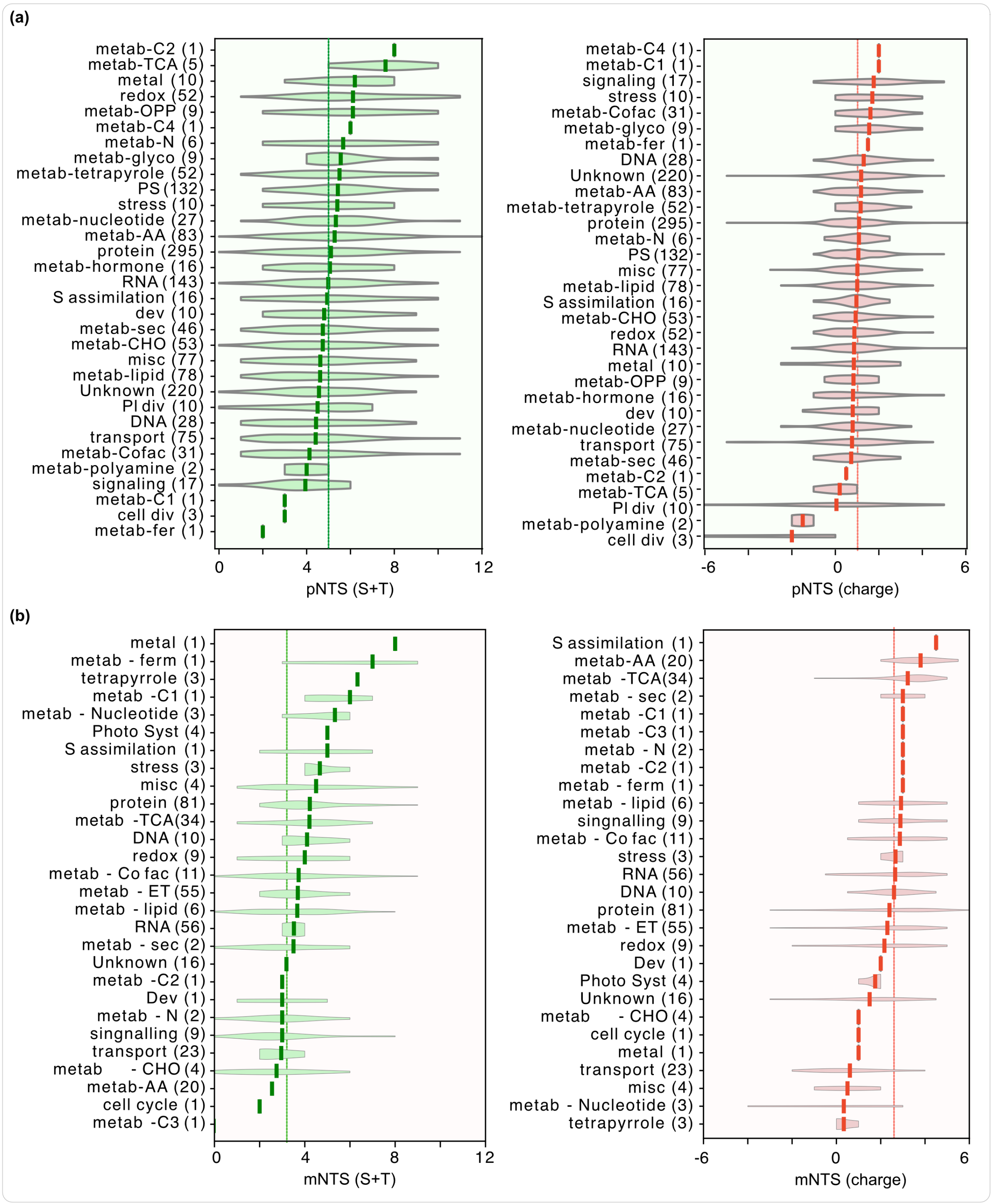


**Fig. S4:** The distribution of the number of phosphorylatable amino acids and charge in the first 20 amino acids of **(a)** plastid and **(b)** mitochondrial proteins in *Arabidopsis*, in each functional category (shown on the left, with number of total proteins in the parenthesis). The vertical lines (running across functional categories) indicate average values for all proteins in that organelle. For full names of the functional categories, please refer to the figure source data.

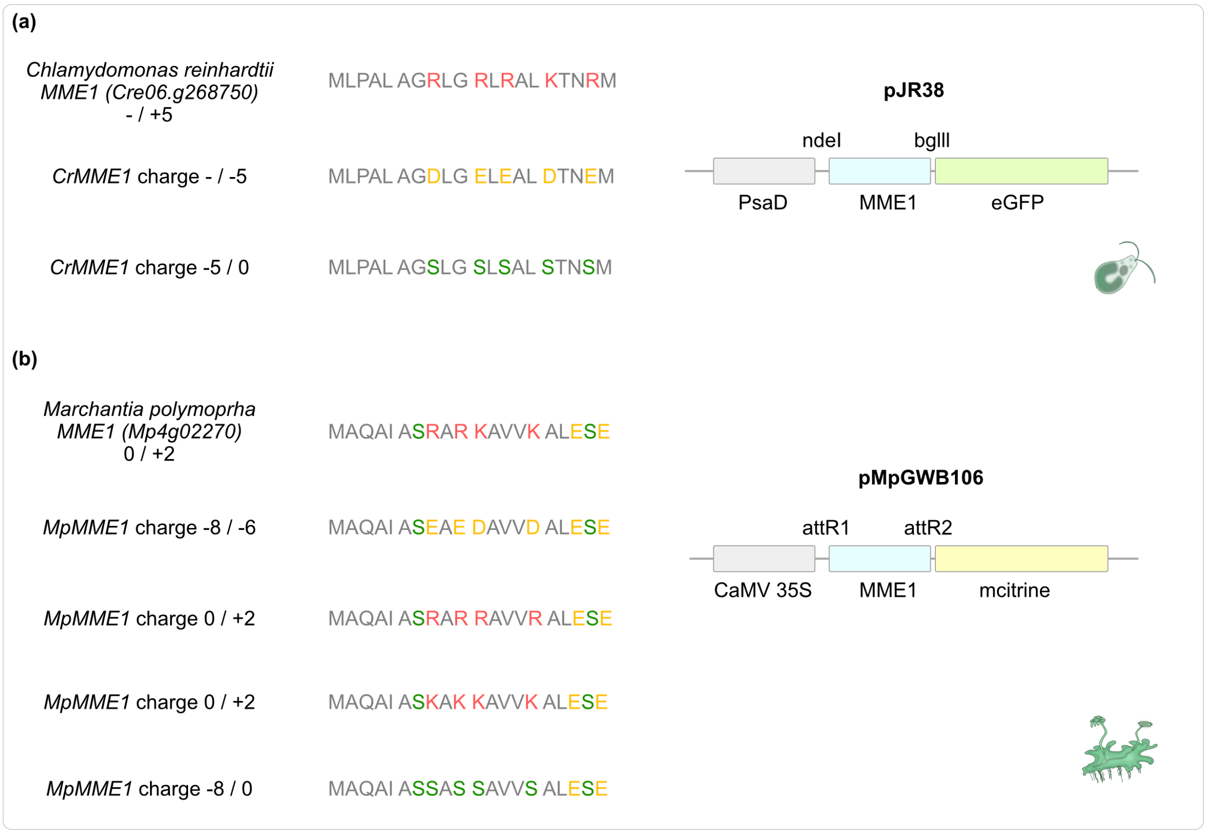


**Fig. S5: Details on the targeting constructs used.** First 20 amino acids of the native (first row for each species) and mutated variants of MME1 in **(a)** *Chlamydomonas* and **(b)** *Marchantia*. Relevant physicochemical amino acids are highlighted in salmon (positively charged), yellow (negatively charged) and green (phosphorylatable). The charges represent the two extremes of phosphorylation, separated by '/', with full (i.e. all serines in the first 20 amino acids are phosphorylated) versus no phosphorylation (i.e. none of the serines in the first 20 amino acids are phosphorylated). The first 20 amino acids of *Chlamydomonas* MME1 were fused at the N terminal of eGFP, under the Photosystem I reaction centre subunit II (PsaD) promoter in vector pJR38. The full length *Marchantia* MME1 was fused at the N terminal of citrine under the Cauliflower mosaic virus promoter.

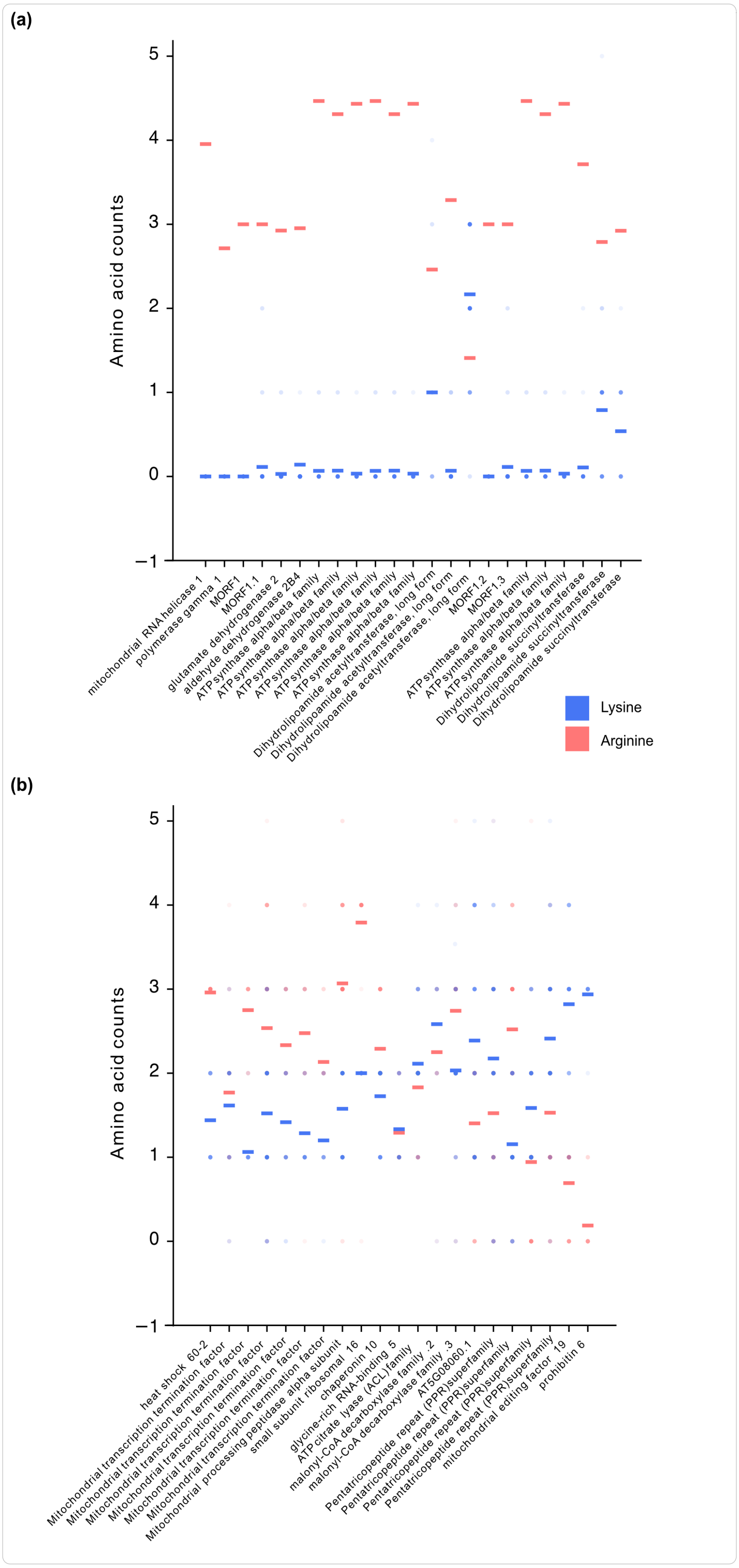


**Fig. S6: Arginine (R) vs. lysine (K) preference across mNTSs of 72 eudicots**. R and K counts from the first 20 amino acids for orthologues of *Arabidopsis* mitochondrial proteins with a charge higher than 3. Proteins showing the strongest preference for **(a)** R and **(b)** K are plotted. Each column is a protein, each data point is a number of R (salmon) and K (blue) in a given species. The horizontal lines indicate the average R (in salmon) and K numbers (in blue) for that protein's orthologues across eudicots.

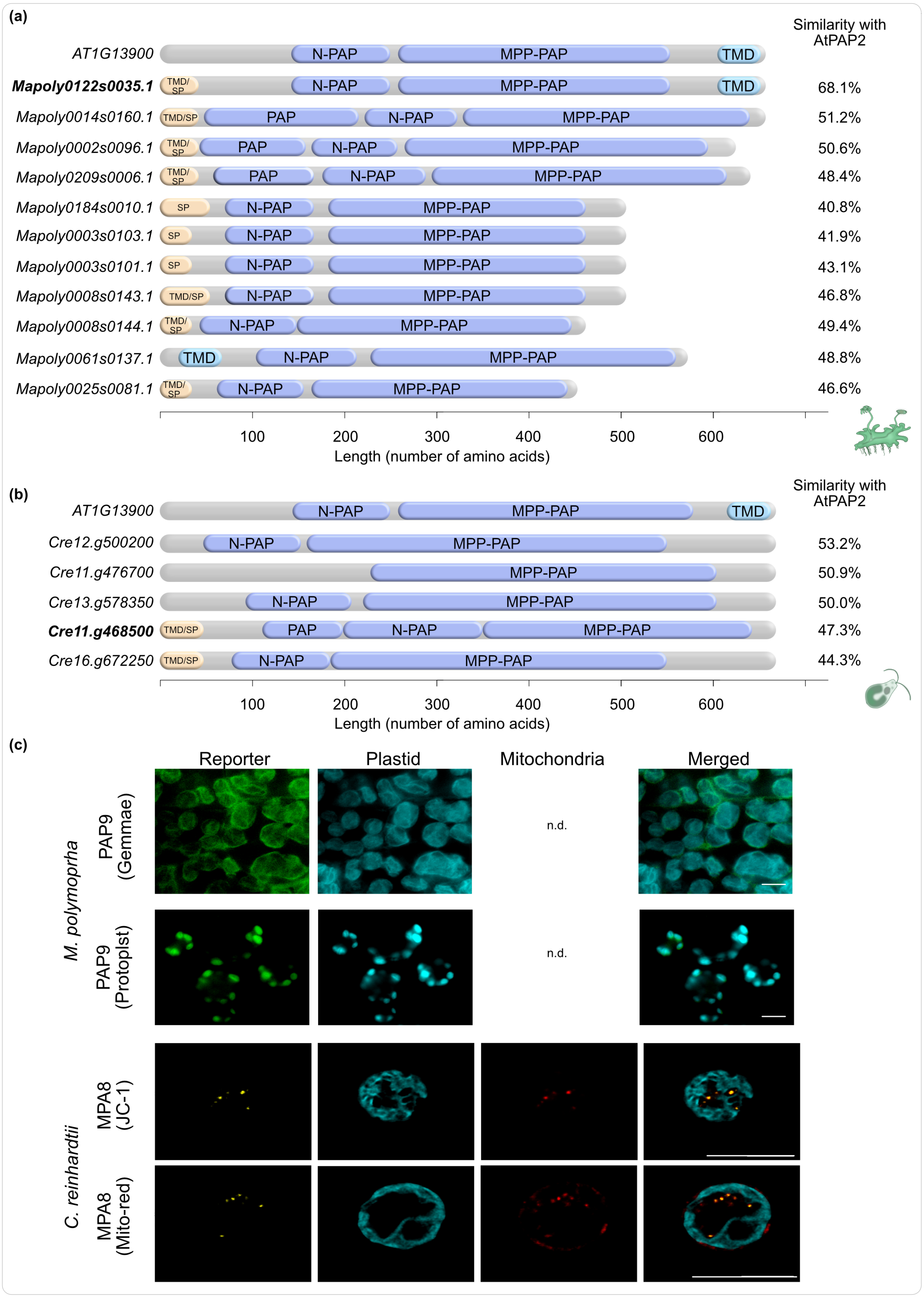


**Fig. S7: Identification and localisation of PAP2/9 homologues in *Chlamydomonas* and *Marchantia*.** *Arabidopsis* PAP2 (AT1G13900) has five homologues in *Chlamydomonas reinhardtii* (genome v5.6). **(a)** Their functional and membrane docking domain analyses narrowed down Cre11.g468500 as the best match, as it has the signature PAP2 domains, as well as a membrane anchoring/signalling peptide with a probability higher than 0.5. The functional domains (N-terminal purple acid phosphatase, N-

PAP; metalloprotein phosphatase, MPP-PAP) were inferred via InterProScan (https://www.ebi.ac.uk/interpro/). Transmembrane domains (TMD) were initially inferred via TMHMM2 (https://services.healthtech.dtu.dk/services/TMHMM-2.0/). The updated version deepTMHMM (https://dtu.biolib.com/DeepTMHMM) annotated the same region in some cases (e.g. the N-terminal region in *Chlamydomonas*) as a signal peptide (SP), and we therefore labelled those regions 'TMD/SP'. Regions with a single label 'TMD' (e.g. at the C-terminus of *At*PAP2) are the ones where TMHMM2 and deepTMHMM both annotated it as TMD. **(b)** Similar to *Chlamydomonas,* all *Marchantia polymoprha* v3.1 homologues were analysed to annotate Mapoly0122s0035 as the best candidate for *Mp*PAP. **(c)** N terminal of *CrMPA8* and the full length MpPAP9 were localized in in the thallus and protoplasts of *Marchantia*, and in cells of *Chlamydomonas.* Plastids were imaged through their autofluorescence and mitochondria through MitoTracker Red and JC-1. Scalebars 5µM.

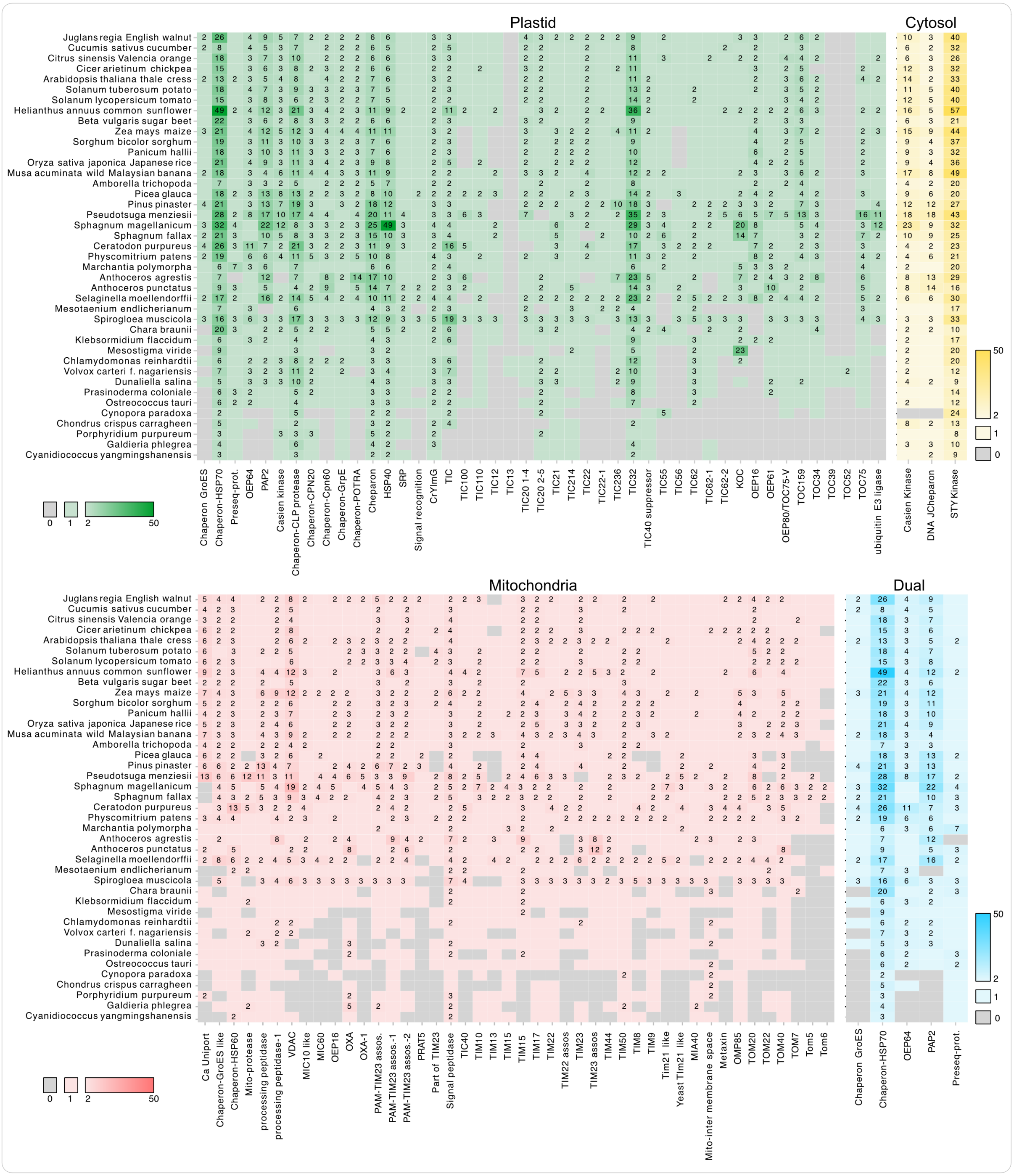


**Fig. S8: Gene copy numbers (GCN) of import components across the Archaeplastida**. Entries are separated based on the intracellular localisation of the components: plastid, cytosol, mitochondrial or dual (plastid and mitochondrial). Colour schemes represent the GCN (components with more than one copy are shown also by the values on each cell).

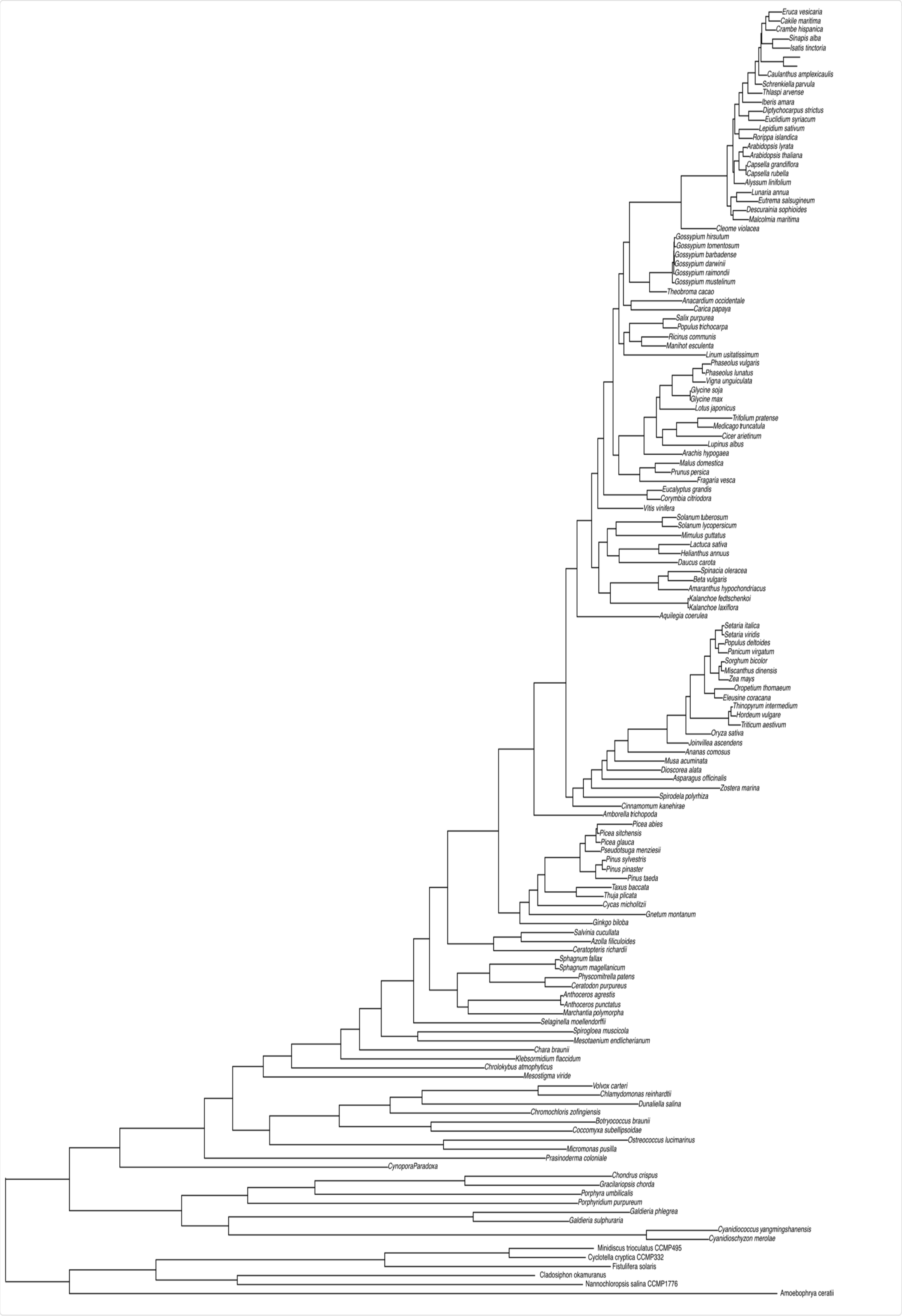


**Fig. S9:** Phylogeny of 137 archaeplastidal species (with six outgroup species).

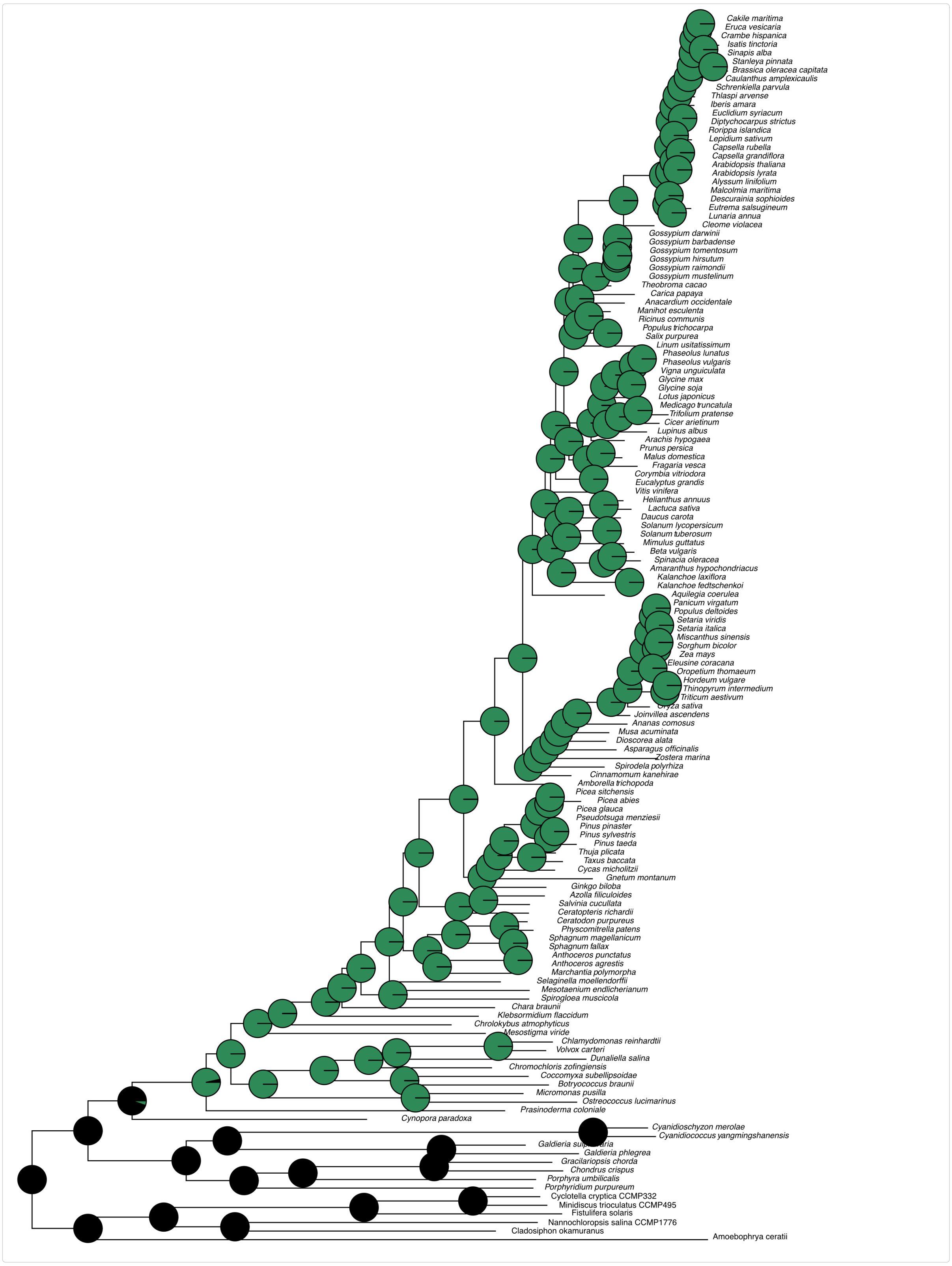


**Fig. S10** Ancestral state reconstruction for TIC56 (the full phylogeny and ASR of the focused snippet in Fig. 5B).

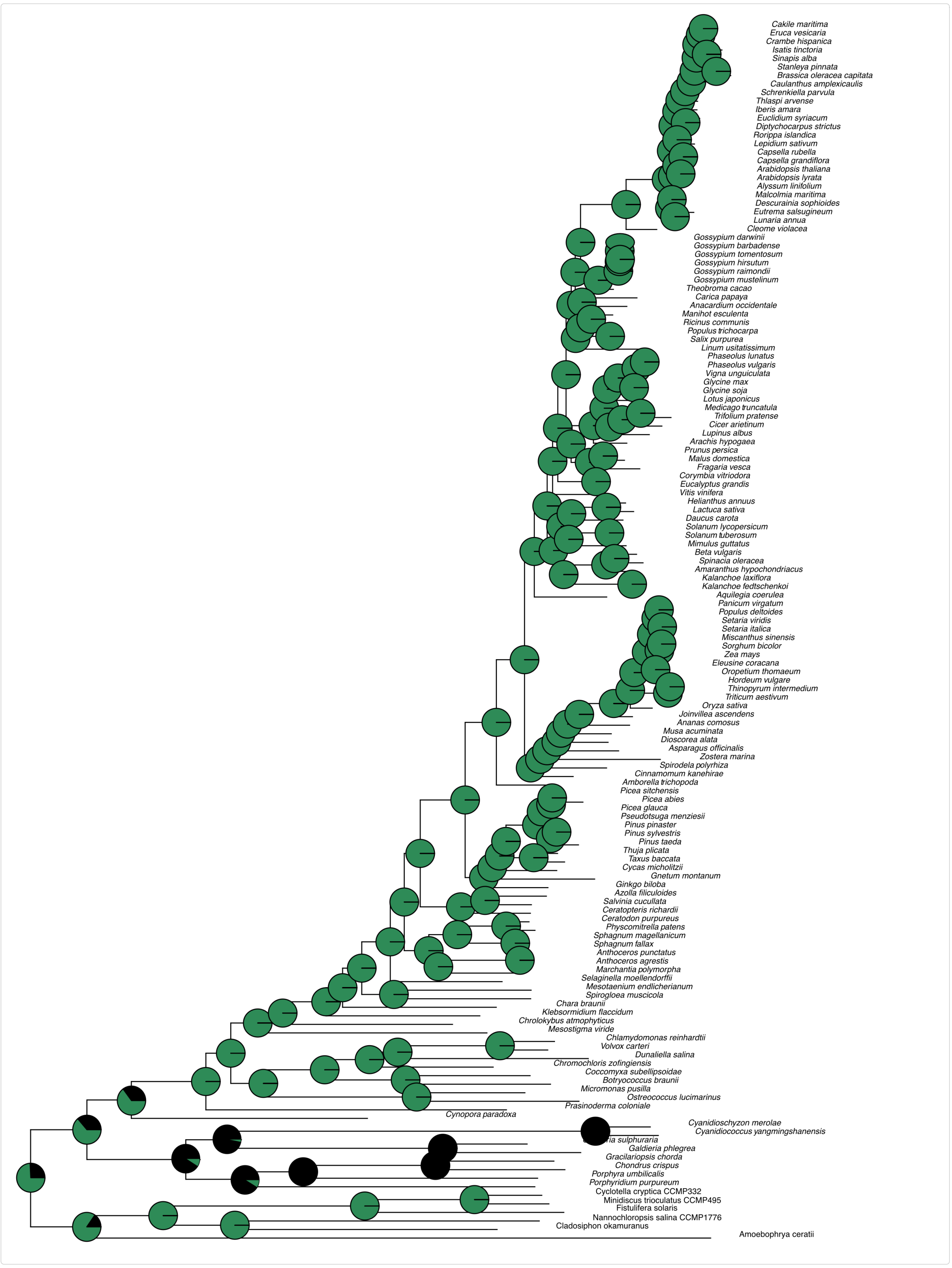


**Fig. S11** Ancestral state reconstruction for TIC100 (the full phylogeny and ASR of the focused snippet in Fig. 5C).

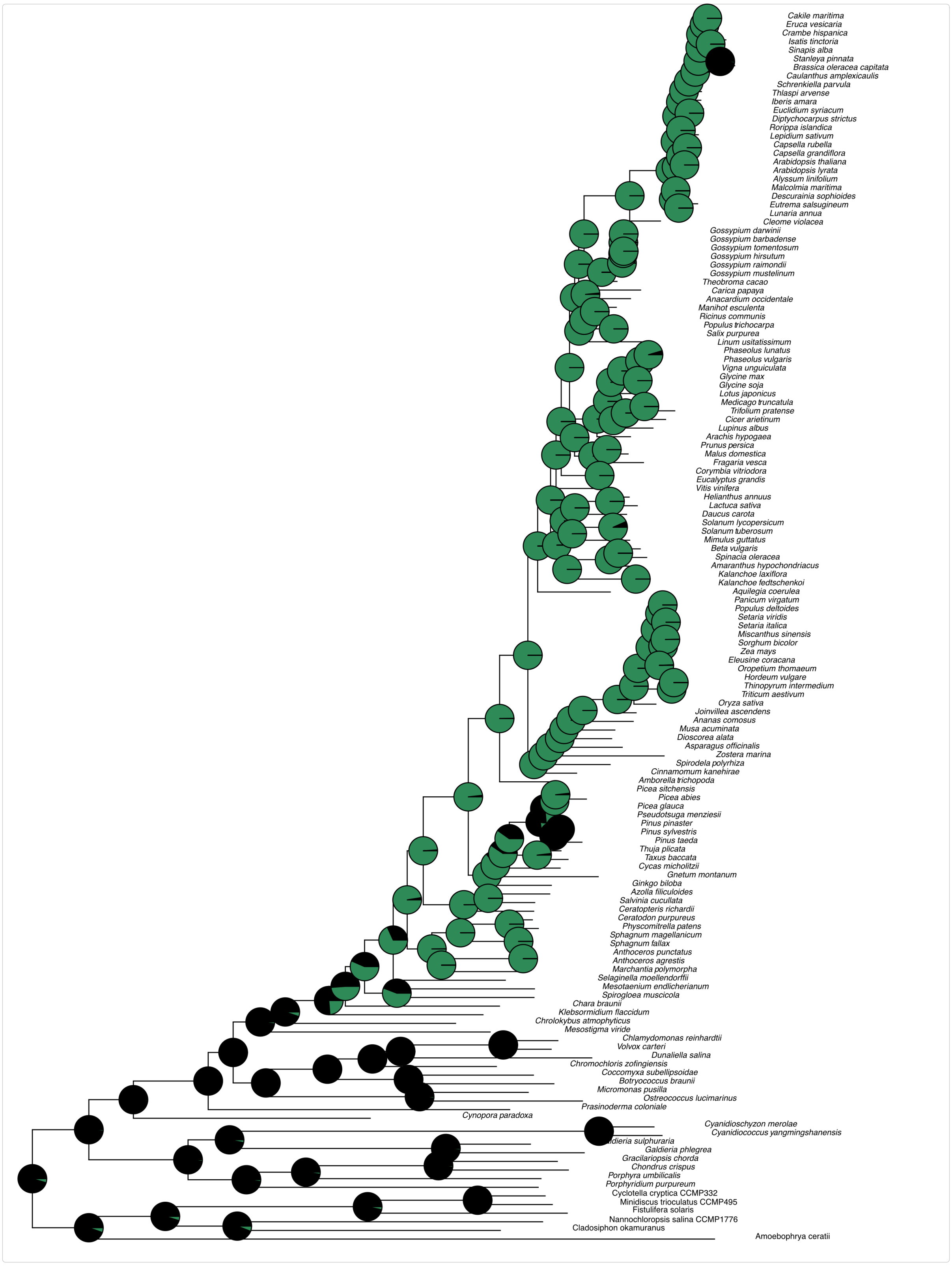


**Fig. S12** Ancestral state reconstruction for TIC12 (the full phylogeny and ASR of the focused snippet in Fig. 5D).

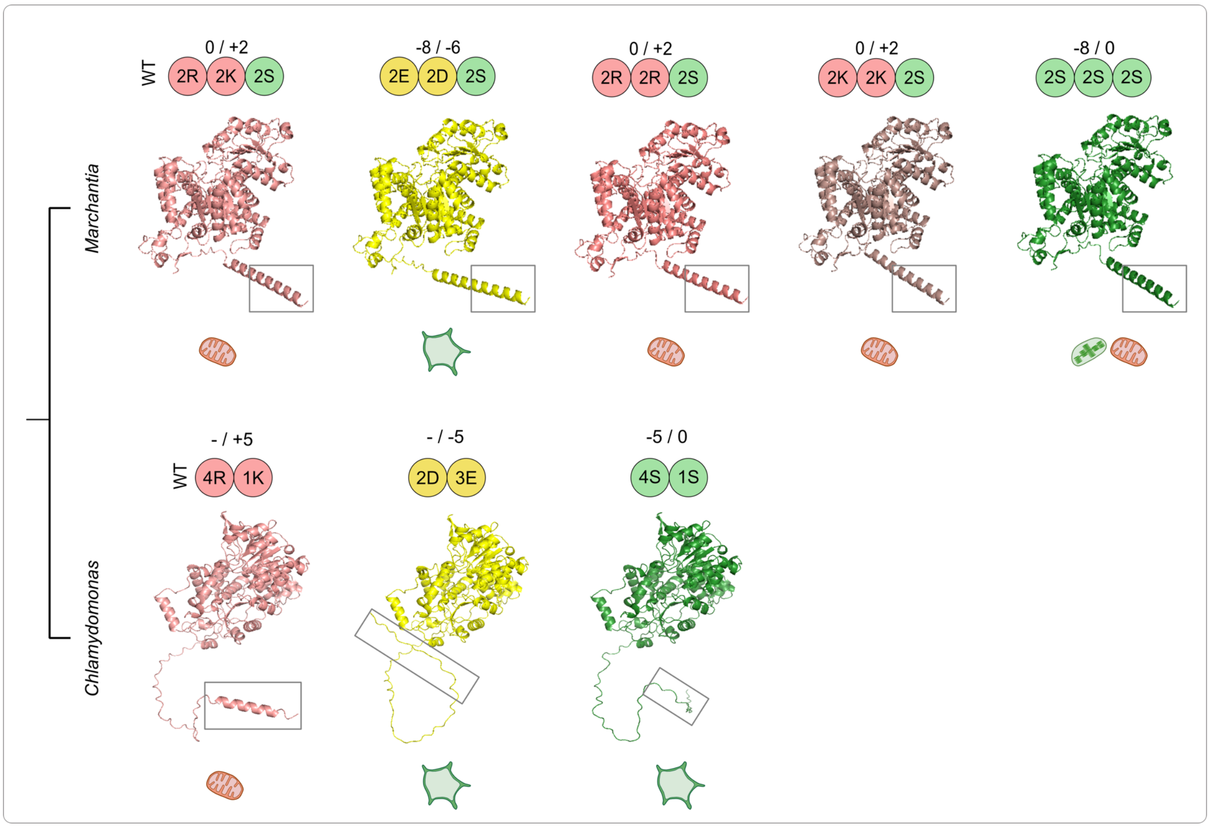


**Fig. S13** Predicted structures of MME1 constructs used in the study and their observed localisation. The key amino acids (in single letter codes) are shown on top of each structure, along with the resulting charge of the first 20 amino acids (with/out phosphorylation). The first 20 amino acids are indicated by a box. The observed localization of each of these constructs is indicated by icons at the bottom of each structure.
